# Shore crabs reveal novel evolutionary attributes of the mushroom body

**DOI:** 10.1101/2020.11.06.371492

**Authors:** Nicholas James Strausfeld, Marcel Ethan Sayre

## Abstract

Neural organization of mushroom bodies is largely consistent across insects, whereas the ancestral ground pattern diverges broadly across crustacean lineages, resulting in successive loss of columns and the acquisition of domed centers retaining ancestral Hebbian-like networks and aminergic connections. We demonstrate here a major departure from this evolutionary trend in Brachyura, the most recent malacostracan lineage. Instead of occupying the rostral surface of the lateral protocerebrum, mushroom body calyces are buried deep within it, with their columns extending outwards to an expansive system of gyri on the brain’s surface. The organization amongst mushroom body neurons reaches extreme elaboration throughout its constituent neuropils. The calyces, columns, and especially the gyri show DC0 immunoreactivity, an indicator of extensive circuits involved in learning and memory.

## Introduction

Insect mushroom bodies, particularly those of *Drosophila*, are the most accessible models for elucidating molecular and computational algorithms underlying learning and memory within genetically and connectomically defined circuits (e.g., Aso et al., 2014; Senapati et al., 2019; Modi et al., 2020; Jacob and Waddell, 2020). An advantage of *Drosophila* is that it exemplifies a ground pattern organization that is largely consistent across hexapod lineages (Ito et al., 1998, 2014; Sinakevitch et al., 2001; Li and Strausfeld, 1997; Montgomery and Ott 2015; Groh and Rössler, 2011).

Because of the importance of mushroom bodies in understanding the relevance of synaptic organization in sentience and cognition, recognizing evolutionary divergence of these centers would be expected to yield experimentally testable predictions about evolved modifications of circuitry in relation to ecological demands imposed on the species. However, insect mushroom bodies show a remarkably conserved organization, which makes them relatively unsuited for neuroevolutionary studies. There are some minor exceptions, to be sure: in honey bees and ants, calycal domains receive modality-specific afferents; in beetles, dietary generalists have more elaborate calyces than specialists; in aquatic beetles a modality switch has replaced olfactory input to a visual input into the calyces (Gronenberg, 2001; Farris and Schulmeister, 2011; Lin and Strausfeld, 2012). Less attention has been given to neural arrangements comprising the mushroom body lobes (columns), although some studies have addressed distinctions in basal groups such as silverfish (Zygentoma), dragonflies and mayflies (Farris, 2005; Strausfeld et al., 2009).

The evolutionary stability of the insect mushroom body contrasts with the recent demonstration that mushroom bodies of malacostracan crustaceans have undergone substantial and often dramatic modification of the mandibulate (ancestral) mushroom body ground pattern (Stegner and Richter, 2011; Wolff et al., 2015, 2017; Strausfeld et al., 2020). Whereas Stomatopoda (mantis shrimps), the sister group of Eumalacostraca, possesses mushroom bodies that correspond to those of insects, equipped with calyces from which arise prominent columns (Wolff et al., 2017), the trend over geological time has been a reduction of those columnar components such that the ancestral ground pattern organization of parallel fibers and orthogonal Hebbian networks has become morphed to provide planar arrangements within domed centers lacking columns (Wolff et al., 2015; Strausfeld and Sayre, 2019; Strausfeld et al., 2020). These neuronal adaptations nevertheless share the property with insect mushroom bodies of being immunopositive to an antibody raised against the catalytic subunit of protein kinase A, encoded by the *Drosophila* gene DC0, that is required for effective learning and memory (Kalderon and Rubin, 1988; Skoulakis et al., 1993). Antibodies raised against DC0 are reliable identifiers of neuropils mediating learning and memory both in arthropods and other phyla (Wolff and Strausfeld, 2012, 2015).

Whereas the evolutionary shift from columnar mushroom bodies to noncolumnar homologues is demonstrated across most decapod crustaceans, one lineage has until now defied unambiguous identification of a columnar or even a noncolumnar center. This is Brachyura, known in the vernacular as true crabs. Brachyura is a comparatively young lineage, recognized from fossils dating from the mid-Jurassic (Schweizer and Feldmann, 2010; Guinot, 2019). Phylogenomics places the origin of Brachyura also as mid-to-late Jurassic (Wolfe et al., 2019). Brachyura is the most species-rich decapod clade, comprising 6,793 currently known species (Ng et al., 2008). It is also a hugely successful lineage, extant species broadly distributed to occupy benthic, littoral, estuarine, brackish, fresh water, terrestrial and even arboreal habitats. Many of these ecologies are defined by complex topographies (Lee, 2015; Hartnell, 1988).

Historically, identifying a mushroom body homologue in the crab’s brain has been problematic. Claims for homologous centers range from paired neuropils in the brain’s second segment, the deutocerebrum, later attributed to the olfactory system (Bethe 1897), to an insistence that the crab’s reniform body is a mushroom body (Maza et al., 2016, 2020). Demonstrated 132 years ago in stomatopod crustaceans, the reniform body is a morphologically distinct center that coexists in the brain’s lateral protocerebrum adjacent to its columnar mushroom bodies (Bellonci, 1882; Thoen et al., 2019).

Until the present, observations of the varunid shore crab *Hemigrapsus nudus* have identified large anti-DC0-reactive domains occupying almost the entire rostral volume of its lateral protocerebrum but no evidence for mushroom bodies (Thoen et al., 2019). The affinity of these domains to anti-DC0 suggested, therefore, a cognitive center far more expansive than found in any other arthropod of comparable size, with the possible exception of the domed mushroom body of the land hermit crab *Coenobita clypeatus* (Wolff et al., 2012). Here we demonstrate that in the shore crab there are indeed paired mushroom bodies. But these have been “hiding in plain sight,” having undergone an entirely unexpected neurological transformation that is opposite to the evolutionary trend shown in other lineages towards a domed noncolumnar morphology.

Here we demonstrate evidence at level of neuronal arrangements showing that in the shore crab paired mushroom bodies have undergone an evolved transformation that is possibly unique to Arthropoda. They are inverted: their calyces reside deep within the lateral protocerebrum, a location allowing outward expansion of the mushroom body columns which reach expanded cortex-like folds beneath the brain’s surface. The entire disposition of the shore crab’s mushroom bodies is opposite to that of any other crustacean, or insect, in which the calyces are situated under the rostral surface of the lateral protocerebrum with their columns extending downwards into deeper neuropil (Strausfeld et al., 2009; Wolff et al., 2017; Sayre and Strausfeld, 2018).

Characters (traits) defining neuronal organization demonstrate that the mushroom bodies of Stomatopoda phenotypically correspond to those defining the mushroom bodies of *Drosophila* (Wolff et al., 2017). The following description of uses complementary methods that resolve those traits in *Hemigrapsus nudus*. Reduced silver staining demonstrates spatial arrangements of calycal and columnar neuropils and their detailed neuroarchitecture. Osmium-ethyl gallate treatment of intact brains resolves neuronal densities; and serial sections of these preparations provide the three-dimensional reconstructions shown throughout this account (see Methods). Golgi mass-impregnations enable crucial insights into the exceptionally elaborate organization of mushroom body intrinsic neurons, identified as the phenotypic homologues of insect Kenyon cells. Immunohistology has been used to resolve DC0-immunopositive components of the mushroom body calyces and columns, and antibodies raised against GAD (glutamic acid decarboxylase) reveal putative levels of local inhibition. Antibodies raised against tyrosine hydroxylase (TH) and 5-hydroxytryptophan (5HT) demonstrate arborizations consistent with those of output and input neurons intersecting different levels of the *Drosophila* and Stomatopod mushroom body columns.

We show that despite the varunid mushroom body’s unparalleled intricacy and its unique transformation of the ancestral ground pattern, its defining traits nevertheless demonstrate phenotypic homology with the mushroom bodies of insects and those of other crustaceans. We close with a Discussion proposing that the evolution of such intricacy, and the expansion of the lateral protocerebrum to accommodate mushroom body enlargement, is likely to have contributed novel circuits and, ultimately, to cognitive flexibility permitting varunid Brachyura to exploit complex ecological topographies at interface of marine and terrestrial biotopes.

## Results

The Results are organized into eight sections, the first describing the overall location and structure of the varunid mushroom body. This is followed by the identification of its neural components and their arrangements as elaborate networks comprising the paired calyces. Descriptions of the internal organization of the calyces’ rostral extensions, and their two prominent columns, culminates with an explanation of the cortex-like organization of overlying gyri to which the columns project. The final observations address the relationship of the mushroom body to the reniform body, a center common to malacostracan crustaceans. To provide a three-dimensional understanding of this highly intricate system, descriptions refer to locations and arrangements within a reconstructed lateral protocerebrum and its major components (*see Methods*).

### 1. The varunid lateral protocerebrum (*Figures 1, 2, Figure 1 - figure supplement 1*)

In crabs, as is typical of stomatopods and most decapod crustaceans, the most anterior part of its brain, the protocerebrum, is flanked each side by a lateral outgrowth that expands into an enlargement of the eyestalk immediately proximal to the compound retina (*Figure 1 - figure supplement 1A*). This volume of brain comprises the lateral protocerebrum (*Figures 1A, 2A*). The lateral protocerebrum is connected to the midbrain proper by the eyestalk nerve composed of axons relaying information to and from the midbrain.

**Figure 1.**
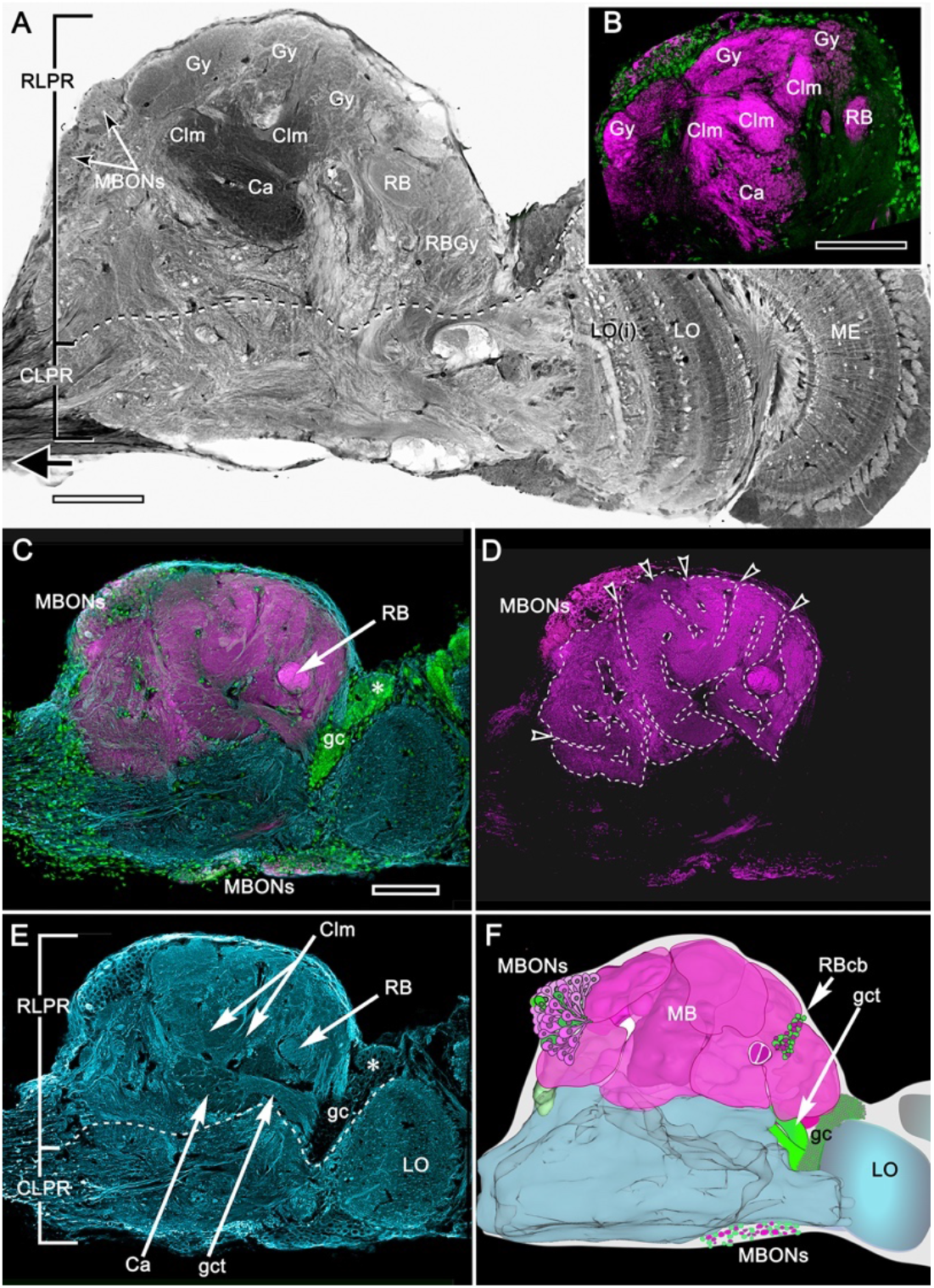
Organization of the varunid crab mushroom body and its associated gyri. (**A**) Brains treated with osmium-ethyl gallate demonstrate the overall disposition of neuropils, exemplified by this section of the lateral protocerebrum from its medial border to the optic lobe medulla. Its rostral domain (RLPR) includes the mushroom body calyces (Ca), columns (Clm) and their associated gyriform neuropils (Gy).Clusters of neuronal cell bodies (here MBONs) are distributed over the neuropil. Distal volumes of the RLPR are associated with the reniform body (RB; here its pedestal and a gyrus RBGy), which is situated between the mushroom body and the optic lobe, here represented by its medulla (ME) and bilayered lobula (LO, LOi). The optic lobe provides outputs to discrete neuropils comprising the caudal lateral protocerebrum (CLPR). (**B-E**) Anti-DC0-labelled (magenta) volumes in panel B match the corresponding membrane-dense calyces and columns and their rostral gyri in panel A. Panel C demonstrates that DC0-immunostained gyri occupy almost the entire rostral level of the RLPR. Two small clusters caudally (MBONs) lining the CLPR, near the optic lobes, also include DC0-immunoreactive perikarya. The nuclear stain Syto13 (green) demonstrates the location of mushroom body globuli cells (gc) lying immediately beneath a group slightly large perikarya (asterisk). In panel D, removal of anti-□ tubulin and Syto13 labelling from C resolves the extent of the gyri and their many sulcus-like indentations (open arrowheads). In panel E anti-□□tubulin shown here alone resolves the overall fibroarchitecture of the LPR to provide correlative data with those provided by reduced silver stains (*Figure 2*). (**F**) Total Amira-generated reconstruction of a serial-sectioned, osmium-ethyl gallate treated eyestalk demonstrates the huge area of the lateral protocerebrum occupied by the folded gyri. Superimposed are the locations of small neurons supplying the reniform body (RBcb), mushroom body globuli cells (gc) and their tract of neurites leading to the calyx (gct), here obscured by overlying gyri. Scale bars, A, B, 100μm; C-F, 100μm. **Figure 1 ––– figure supplement 1.** Schematic of the *Hemigrapsus* brain

**Figure 2.**
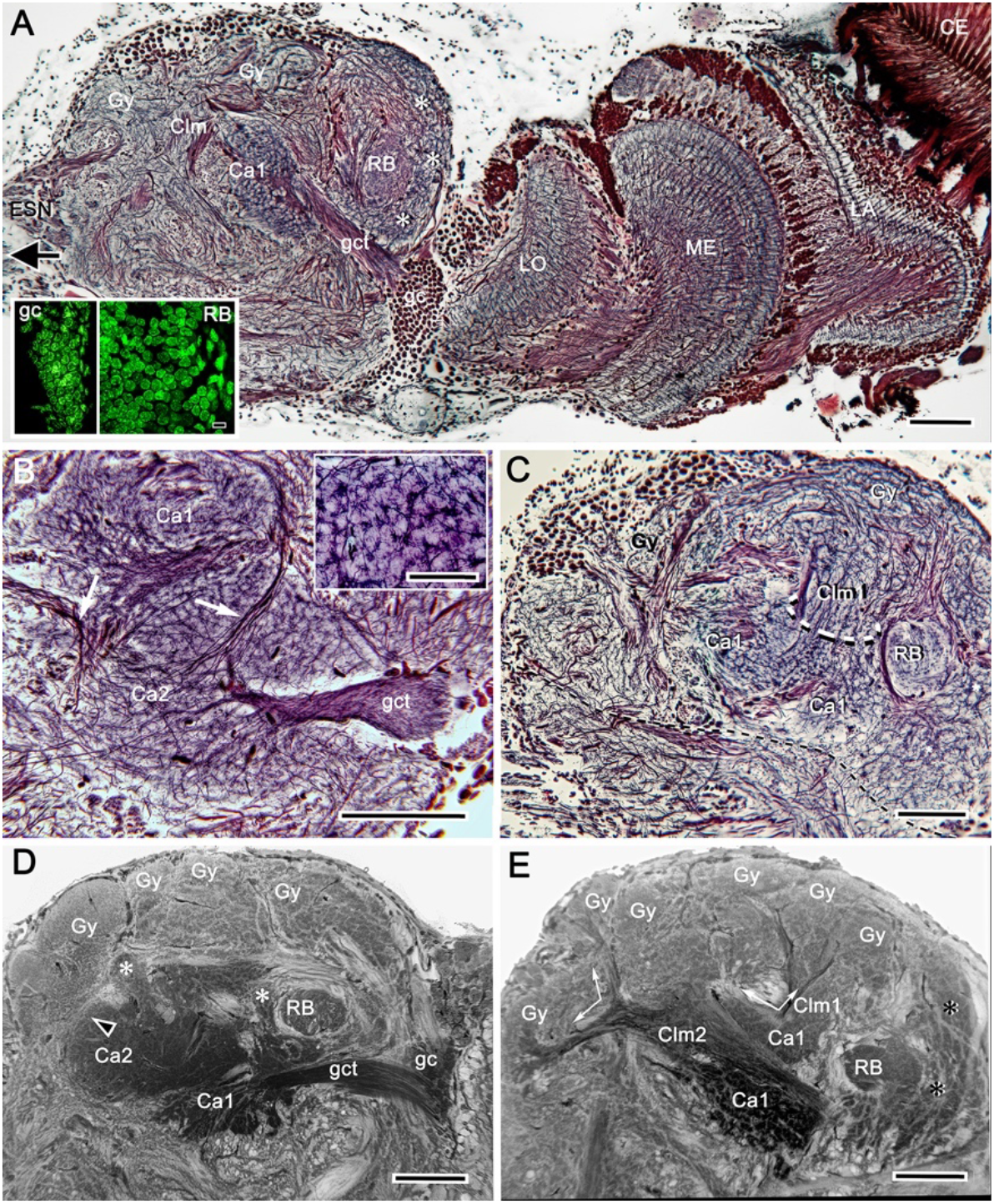
Uniquely identifiable neuropils of the mushroom bodies are resolved by reduced silver. (**A**) Horizontal section of the lateral protocerebrum demonstrating the more dorsal of the two calyces (Ca1) and its supply by the globuli cell tract (gct) from the cluster of globuli cells (gc). Columnar extensions (Clm) from the calyces reach out to overlying gyri (Gy). The reniform body is shown with three of its gyri (asterisks) and its bundled axons (RB) situated between the mushroom body and optic lobes, insets lower left compare the smallest perikarya supplying the calyces (globuli cells, gc) with the next smallest perikarya belonging to neurons of the reniform body (see *Figure 2 -figure supplement 1*). Abbreviations: ESN, eyestalk nerve; CE, compound retina, LA, lamina, ME, medulla; LO, lobula. (**B**) Adjacent calyces (Ca1, Ca2) showing their diagnostic arrangement of microglomeruli (inset upper right; see also Figure 3). Arrows indicate afferent fibers into Ca2 from nonolfactory origins. (**C**) The calycal origin and rostral extension of the Ca1 column (encircled). At this level, penetration by en passant axon bundles causes the calyx to appear fragmented. (**D, E**) Osmium ethyl gallate-treated sections at two depths. Panel D indicates the bulk and elaboration of calycal neuropils. At this level calyx 2 appears as the larger. The arrowhead indicates the root of column 2. Calyx 1 is shown with its globuli cell tract entering a domain denoted by lateral extensions of calyx 1. In panel E, the base of column 1 (Clm1) is shown arising from Ca1, its curvature at this level splitting it into two parts. Column (Clm2) is shown extending two branches into the gyriform layer above. Clm1 extends outwards just proximal to the pedestal of the reniform body (RB). In addition to columns, calyces provide occasional narrower finger-like extensions extending towards the gyri (asterisks in D). Scale bars, 100μm. Inset in A, 10μm, inset in B, 50μm. **Figure 2 ––– figure supplement 1.** Globuli cells supplying intrinsic neuron to the mushroom bodies **Figure 2 ––– figure supplement 2.** Globuli cells provide two tracts, one to each calyx.

Brains treated with osmium-ethyl gallate or with reduced silver (*Figures 1A, 2A*) resolve the varunid lateral protocerebrum as a hump-shaped volume divided into a rostral and caudal part. The rostral lateral protocerebrum (RLPR), once referred to as the terminal medulla, a now defunct term (Strausfeld, 2020), is dominated by two centers: the mushroom bodies and the smaller reniform body. There is a small number of other satellite neuropils near the origin of the eyestalk nerve. The mushroom body is immediately recognizable by its intensely osmiophilic calyces from which arise columns that extend outwards, branching into a cortex-like neuropil composed of gyri (*Figure 1A*). These occupy the outermost volume of the RLPR. The reniform body comprises an independent system of interconnected neuropils situated laterally constrained to a volume of the RLPR that extends from its dorsal to its ventral surface (Thoen et al., 2019). In longitudinal sections of the RLPR, its prominent axon bundle, the pedestal, is seen in cross section(*Figures 1A, 2A*). The crab’s RLPR corresponds to that volume of the insect protocerebrum containing the mushroom body and its immediate associated neuropils, such as the lateral horn (*Figure 1 - figure supplement 1B, C*).

As in stomatopods and decapods, the crab’s RLPR receives terminals of olfactory projection neurons originating from both the ipsi- and contralateral olfactory (antennular) lobes located in the brain’s second segment, the deutocerebrum (*Figure 1 - figure supplement 1A*). The axons of more than a thousand projection neurons are organized into several discrete fascicles bundled together to provide the antennoglomerular tract (AGT). The left and right AGTs extend superficially across the midbrain to meet just dorsal to the central body. There, each tract bifurcates to provide inputs to each lateral protocerebrum from both the ipsi- and contralateral olfactory lobes (*Figure 1 - figure supplement 1A*). The eyestalk nerve also carries other efferent axons to the lateral protocerebrum: these include relay neurons from other sensory neuropils, relays from the opposite lateral protocerebrum and optic lobes, and interneurons relaying from various midbrain neuropils.

The caudal volume of the lateral protocerebrum (CLPR) is mainly associated with outputs from the optic lobe neuropils (*Figure 1A*). Most of these originate from the lobula, each bundle of outgoing axons representing a population of one of the numerous morphological types of columnar neurons. Bundles segregate to a system of interconnected neuropils called optic glomeruli, as occurs in insects (Okamura and Strausfeld, 2007). Further relays from the glomeruli send information centrally to the midbrain via the eyestalk nerve. The nerve also carries to the midbrain the axons of neurons from the lateral protocerebrum’s rostral neuropil, the mushroom bodies, and its associated gyri and from CLPR neuropils associated with the optic glomeruli.

### 2. Organization of DC0-positive volumes (Figure 1)

The dense osmiophilic calyces and their extended columns (*Figure 1A*) correspond to discrete neuropils showing high levels of immunoreactivity to antibodies raised against DC0 (*Figure 1B*). The fibroarchitecture of these DC0-positive volumes is revealed by labeling with antibodies against α-tubulin or by reduced silver. (*Figures 1E; 2A-C*) that provide correlative data for resolving and interpreting the arrangements of the mushroom body volumes in 3-D reconstructions (*Figure 2 - figure supplement 2*: see Methods). Salient both in osmium–ethyl gallate-stained brains is the layer of gyri that defines the outer level of the rostral lateral protocerebrum (*Figures 1A, 2D, E*). Gyri are DC0-immunoreactive and are contiguous with each of the DC0-immunoreactive columns that extend outwards from the calyces. Gyri are resolved as pillowed folds and indentations (sulci: *Figure 1D, F*) comparable to the folded architecture of a mammalian cerebral cortex. This feature appears to be unique to Brachyura (see Discussion).

### 3. The calyces and their origin from globuli cells (Figures 1, 2)

If centers are claimed as phenotypic homologues of insect mushroom bodies, then the centers are expected to have the following categories of neurons irrespective of their divergence from the ancestral ground pattern as represented in the allotriocarid clade Cephalocarida (Stegner and Richter, 2011). These categories are: 1) intrinsic neurons, the most populous of which are Kenyon cells defining the mushroom body’s volume, shape, and subdivisions; 2) extrinsic neurons providing inputs from sensory centers (Kanzaki et al., 1989; Lin and Strausfeld, 2012; Li et al., 2020), as well as protocerebral regions encoding high level multisensory information (Li and Strausfeld, 1999); and 3) output neurons that encode information computed by levels of the mushroom body lobes (Li and Strausfeld, 1999; Turner et al., 2008). The latter are referred to here as MBONs, adopting the term from studies on *Drosophila* (Aso et al., 2014a,b).

Intrinsic neurons supplying the *Hemigrapsus nudus* calyces correspond to insect Kenyon cells. As do insect Kenyon cells, these originate from the brain’s smallest and most crowded population of neuronal cell bodies, the globuli cells (*Figures 1C, 2A,D, Figure 2 – figure supplement 1*). In other arthropods globuli cell perikarya have been shown as dense clusters situated at the rostral surface of the lateral protocerebrum, usually near its proximal margin or immediately above the mushroom body calyces, as in *Drosophila* or Stomatopoda, or, in malacostracans with eyestalks, close to the lateral protocerebrum’s attachment to the eyestalk nerve. In *H. nudus*, however, globuli cell perikarya are found in a completely different and surprising locality: a densely populated “hidden” volume tucked between the proximal margin of the lobula and the most distal border of the lateral protocerebrum (*Figure 2A*). As in other pancrustaceans these perikarya are the smallest perikarya of the brain, in the varunid ranging from 5 to 7 μm in diameter (compared in *Figure 2A, inset, Figure 2 – figure supplement 1*). In crabs used for these studies (carapace widths of 4-8 cm), we estimate that this perikaryal cluster accounts for 22,000 mushroom body intrinsic neurons shared by the two calyces (see Methods). Cytological distinctions amongst cells in the cluster, denoted by their appearance after osmium-ethyl gallate staining (Wigglesworth, 1957), suggest that new globuli cells are being continuously generated from ganglion mother cells situated at the cluster’s periphery (*Figure 2 - figure supplement 1D-G*).

Cell body fibers (neurites) extending from the globuli cell cluster are tightly packed into two prominent bundles, called globuli cell tracts, one projecting obliquely ventrally the other obliquely dorsally to supply each of the two adjacent calyces (*Figure 2A, B, D - figure supplement 2*). Neurites comprising these tracts fan out to supply each calyx with its ensembles of mostly orthogonally arranged dendritic networks that interweave amongst a mosaic of microglomeruli (*Figure 2B; Figure 3A, B, - figure supplement 1*). The networks, which are resolved by silver stains, comprise overlapping populations of intrinsic neuron processes extending amongst a mosaic of many thousands of discrete microglomeruli (*Figures 2B, 3A*). These are assumed to be sites of synaptic convergence (*Figure 3-figure supplement 1*), corresponding to microglomerular organization in the calyces of insect, stomatopod and caridean mushroom bodies (Yasuyama et al., 2002; Sjöholm et al., 2006; Groh and Rössler, 2011; Wolff et al., 2017; Sayre and Strausfeld, 2019). As will be described in the following section, components of microglomeruli include the terminals of afferent relays originating from the olfactory lobes, neuropils associated with the visual system, including the reniform body, and centrifugal neurons from the midbrain.

**Figure 3.**
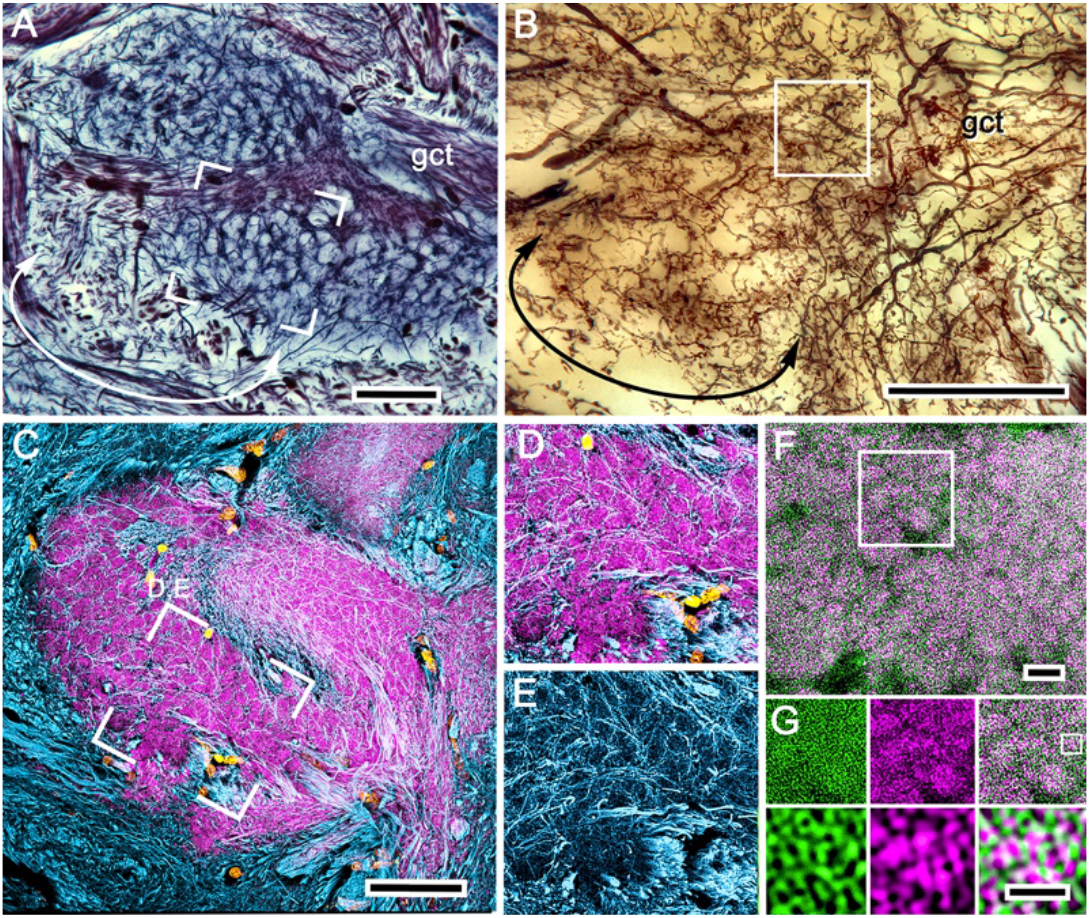
Organization of calyx 1: microglomeruli. (**A**) Reduced-silver-stained calyx showing its globuli cell tract (gct) with, at a different level, many of its constituent neurites giving rise to branches that delineate the mosaic organization of microglomeruli. (**B**) Golgi impregnations provide examples of intrinsic neurons, the branches and fine processes of which extend through domains of the calyx that correspond to those revealed both by reduced silver and by antibodies against DC0. A subfield (curved double arrow) in panel A corresponds to the denoted branches of the illustrated intrinsic neuron in panel B. (**C**) Microglomeruli are specifically labelled by anti-DC0. Microglomerular size and density in the boxed area corresponds to an equivalent microglomerular organization identified by reduced silver (boxed areas in A). (**D-G**) The mosaic is also reflected by the pattern of all intrinsic neuron processes at that level revealed by anti-α-tubulin; these enwrap DC0-positive microglomeruli (D, E). The same system of microglomeruli is resolved using combined actin/anti-synapsin. The area identified in a field of Golgi-impregnated intrinsic neuron processes (boxed in B) is equivalent to the area of the calyx in panel F labelled with actin/anti-synapsin demonstrating the microglomerular mosaic. The boxed area in F is shown in panel G. (**G**) The upper row shows actin and synapsin separate, with the superimposed images in the right panel in G). The boxed area in the upper right panel of G indicates a single microglomerulus. This is enlarged in the lower three panels of G, again showing actin and synapsin as separate images and then combined lower right to demonstrate the density of synapsin-labelled profiles, and thus direct evidence of massive convergence at a calycal microglomerulus (panel G). Scale bars, A–C, 50μm; F, 10μm; G, 2μm. **Figure 3 ––– figure supplement 1.** Rectilinear network organization of the calyx and afferent supply.

Serial sections oriented horizontally, approximately parallel to the rostral surface of the protocerebrum and stained with reduced silver, suggest the calyces are planar neuropils (*Figure 2B*). But this is deceptive, as shown by sections in the vertical plane demonstrating how the deep calycal cytoarchitecture extends rostrally to merge with the base of its column(*Figure 2C*). The total extent and volume of calycal neuropil is surprisingly large as can be seen in intact brains treated with osmium-ethyl gallate, a technique that selectively colors cell membranes black (Wigglesworth, 1957). The method reveals volumes of neuropil in shades of black to light gray, the darker the neuropil, the more densely packed its neuronal processes (*Figures 1A, 2D, E; Figure 4A, figure supplement 1*). This density is a reflection of the many thousands of intrinsic neurons comprising the calyces. Volumes that are paler are composed of neuronal processes that are stouter and contribute less stainable membrane, as in the overlying gyri.

Computer-generated reconstructions and rotations of osmium-ethyl gallate stained sections (see Methods) demonstrate that the calycal volumes are topographically elaborate (*Figure 2 supplement 2*); each calyx is stratified at deeper levels of the lateral protocerebrum’s rostral volume, but where calycal neuropils extend rostrally their upper borders are defined by irregular arrangements of small peaks and troughs. Two prominent columns, one from each calyx, extend outwards into the overlying gyri (*Figure 2E*). Reduced silver material and Golgi impregnations resolve their elaborate internal organization (*Figure 2C*), which is described in section 5, below.

### 4. Neuronal organization of the calyces (Figures 3-6)

Intrinsic neuron processes and terminals of efferent neurons are the main constituents of the varunid calyces (*Figure 3B, Figure 3 - figure supplement 1; Figure 4B-G, Figure 4 - figure supplement 1*). They receive inputs from neurons primarily from the deutocerebrum (*Figure 3 – figure supplement 1B*) but also from the optic lobes and from its associated neuropils, such as the reniform body (*Figure 4 - figure supplement 2*), as also shown for Stomatopoda (Thoen et al., 2019). Golgi impregnations and osmium-ethyl gallate demonstrate that calyces, across their horizontal and vertical extent, are composed of discrete domains of morphologically distinct networks comparable to the discrete modality-specific zones of Kenyon cell dendrites that define the insect calyces (Ehmer and Gronenberg, 2002; Strausfeld, 2002).

**Figure 4.**
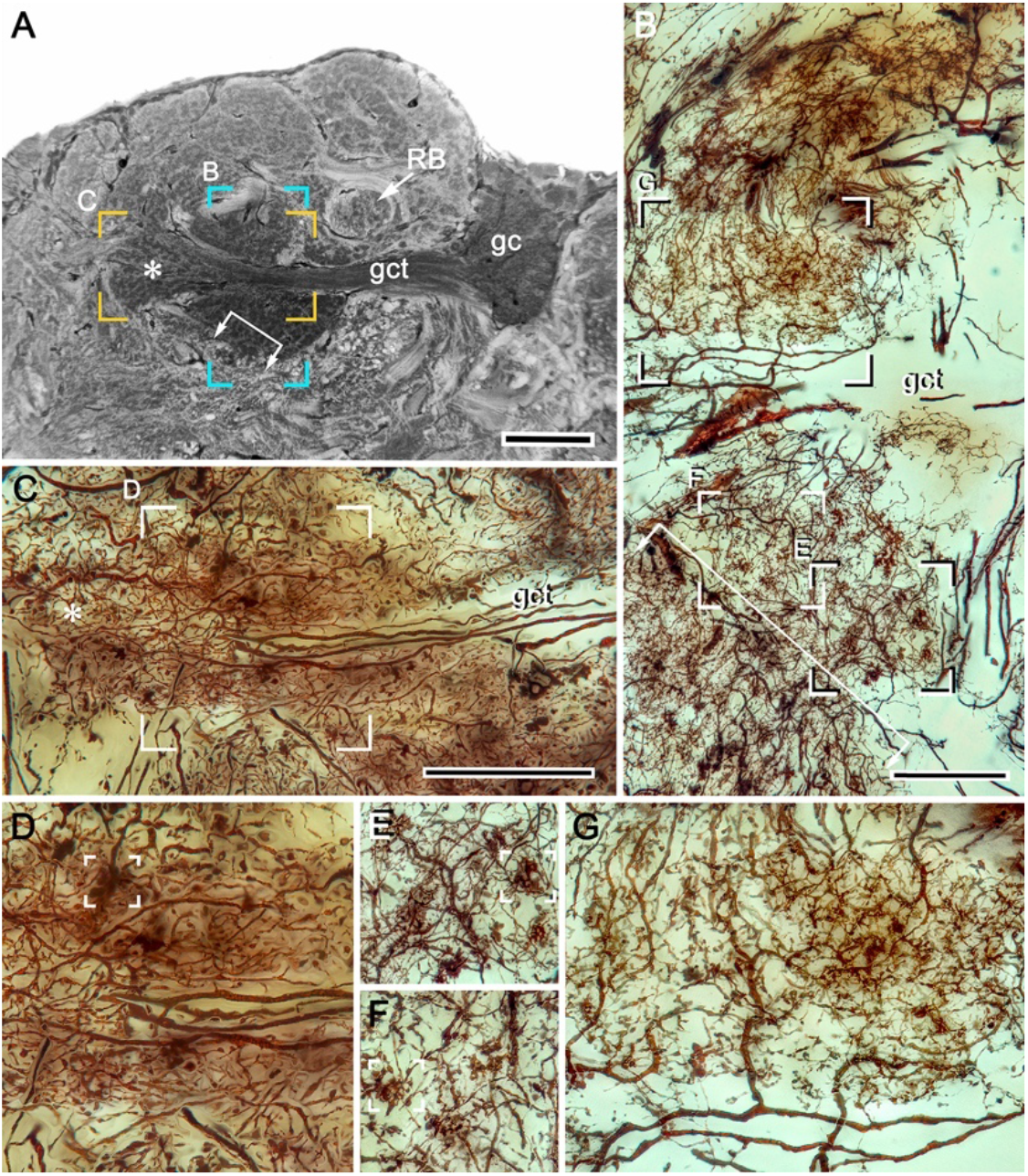
Intrinsic cell dendrites define calycal territories. (**A**) Osmium-ethyl gallate resolves calycal territories matching the dendritic fields of morphologically distinct intrinsic neurons. Letters in panel A indicate panels showing corresponding locations of Golgi-impregnate dendritic fields shown in B and C. Boxed areas in B and C are enlarged in panels D–G to demonstrate distinctive dendritic morphologies. The locations of the arrowed bracket and asterisk in A correspond to their locations in B (the bracket) and C (the asterisk). These further indicate the fidelity of calyx organization indicated by these two histological methods. Open boxes in panels D–-F indicate different morphological specializations of afferent terminals. Abbreviations: RB, reniform pedestal; gc, globuli cells; gct, globuli cell tract. Scale bars, A, C, 100μm, B, 50μm. **Figure 4 ––– figure supplement 1.** Fidelity of intrinsic neurons fields **Figure 4 ––– figure supplement 2.** Calyx distinctions

Most calycal neuropil is characterized by its dense arrangement of small almond-shaped subunits (*Figure 3A*), here referred to as microglomeruli following the term given to comparable elements described from insect mushroom body calyces (Yasuyama et al., 2002 Sjöholm et al., 2006; Groh and Rössler, 2011). However, some peripheral volumes of the calyces comprise more distributed arrangements (*Figure 3A,B*). These appear to receive terminals from axons of relay neurons emerging from fascicles in the eyestalk nerve that are not part of projections from the olfactory lobes (*Figure 2,B)*.

Discrete fields comprising intrinsic cell processes are recognized by their distinctive and particular morphologies (*Figure 4B–G, Figure 4 – figure supplement 1*). Each morphological type is defined by the patterning of its initial branches from its globuli cell neurite and the subsequent reticular trajectories made by their tributaries down to the details of their final specializations. The latter indicate points of functional connection and are morphologically comparable to the claws, clasps and dendritic spines that decorate the dendritic branches of Kenyon cells in *Drosophila* and other insects (Ito et al., 1998; Strausfeld and Li, 1999; Strausfeld 2002) and the dendrites of homologous intrinsic neurons in the mushroom bodies of marine hermit crabs (Strausfeld and Sayre, 2019). As in those taxa, in the varunid calyx these specializations invariably relate to the distribution and patterning of its microglomeruli (*Figure 3A,B*).

Microglomeruli revealed by reduced silver beautifully match the size and mosaic arrangement of microglomeruli resolved by their intense immunoreactivity to antibodies raised against DC0 (*Figure 3 A–C*). The bundled branches of intrinsic neurons resolved by antibodies against α-tubulin (*Figure 3D,E*) further resolve microglomeruli as participants of an elaborate organization of orthogonal networks that correspond to distinct morphological types of intrinsic neurons revealed as single cells by Golgi-impregnation (*Figure 4B,C, Figure 4 - figure supplement 1*). Each glomerulus signifies the site of an incoming terminal’s presynaptic bouton or varicosity (*Figure 3 - figure supplement 1*). Combined anti-synapsin immunoreactivity and F-actin labelling (*Figure 3F-G*) further demonstrates the profusion of postsynaptic sites belonging to intrinsic neurons that converge at each microglomerulus. The densely packed presynaptic specializations further confirm the status of a microglomerulus as the site of massive transfer of information from presynaptic terminals onto many postsynaptic channels.

As remarked, the shapes and arrangements of intrinsic neurons and their synaptic specializations contribute to orthogonal networks in the calyces and define calycal domains. However, the domains are not arranged as neatly adjacent territories as in the insect calyx (Gronenberg, 2001). Instead, the varunid calyx can reveal domains that extend vertically or horizontally (*Figure 3 - figure supplement 1*), or comprise unexpectedly elaborate arrangements that are understood only after they are matched to an appropriately oriented osmium-ethyl gallate section (*Figure 4A,B, figure supplement 1*). One of the most challenging of these arrangements consists of a population of intrinsic neurons defining a cylindrical domain that splays out as it extends towards the proximal margin of the lateral protocerebrum (*Figure 4A*). The entire ensemble comprises densely branched narrow-field arborizations that surround the initial branches of a subset of globuli cell neurites (*Figure 4A,C,D*). The core leads to a terminal expansion that is confluent with the base of one of the two large columns that extends to the overlying gyri. Serial reduced-silver sections reveal this as originating only from calyx 2, indicating that calyx 2 differs from calyx 1 with regard to at least one population of intrinsic neurons. As in many insect lineages where there are paired calyces, it is such subtle distinctions that demonstrate disparate organization across paired calyces (Farris and Strausfeld, 2003). Reduced-silver stains showing axon tracts reveal inputs to calyx 2 originating from a tract that recruits its axons from the reniform body and the medulla. Collaterals from this tract spread into a deep level of calyx2 neuropil where they spread across microglomeruli bordering one side of the incoming neurites from the globuli cell tract (*Figure 4 - figure supplement 2*). The correspondence of this arrangement with insect calyces is considered in the Discussion.

There is one critical exception to the generalization that calycal subdivisions are denoted by different types of intrinsic neurons. The exception is that there is a system of neurons that arborizes uniformly throughout both calyces. These arborizations belong to GAD-immunoreactive local processes originating from a dorsal cluster of large cell bodies, each of which provides an anaxonal field that arborizes throughout both calyces, and into each of its domains. (*Figure 5A,B*). These densely packed processes are exceptional in that they do not obey the orthogonal leitmotif of the calyx, as revealed by anti-a-tubulin (*Figure 5C, D*), nor do they reflect the mosaic arrangement of glomeruli (Figure *5C-E*). Instead, anti-GAD-immunoreactive branchlets populate all the calycal domains and do so in great numbers, the totality suggesting that local inhibitory elements are distributed amongst all intrinsic cell processes irrespective of whether they are in immediate proximity to or distant from a microglomerulus (*Figure 5 C-E*).

**Figure 5.**
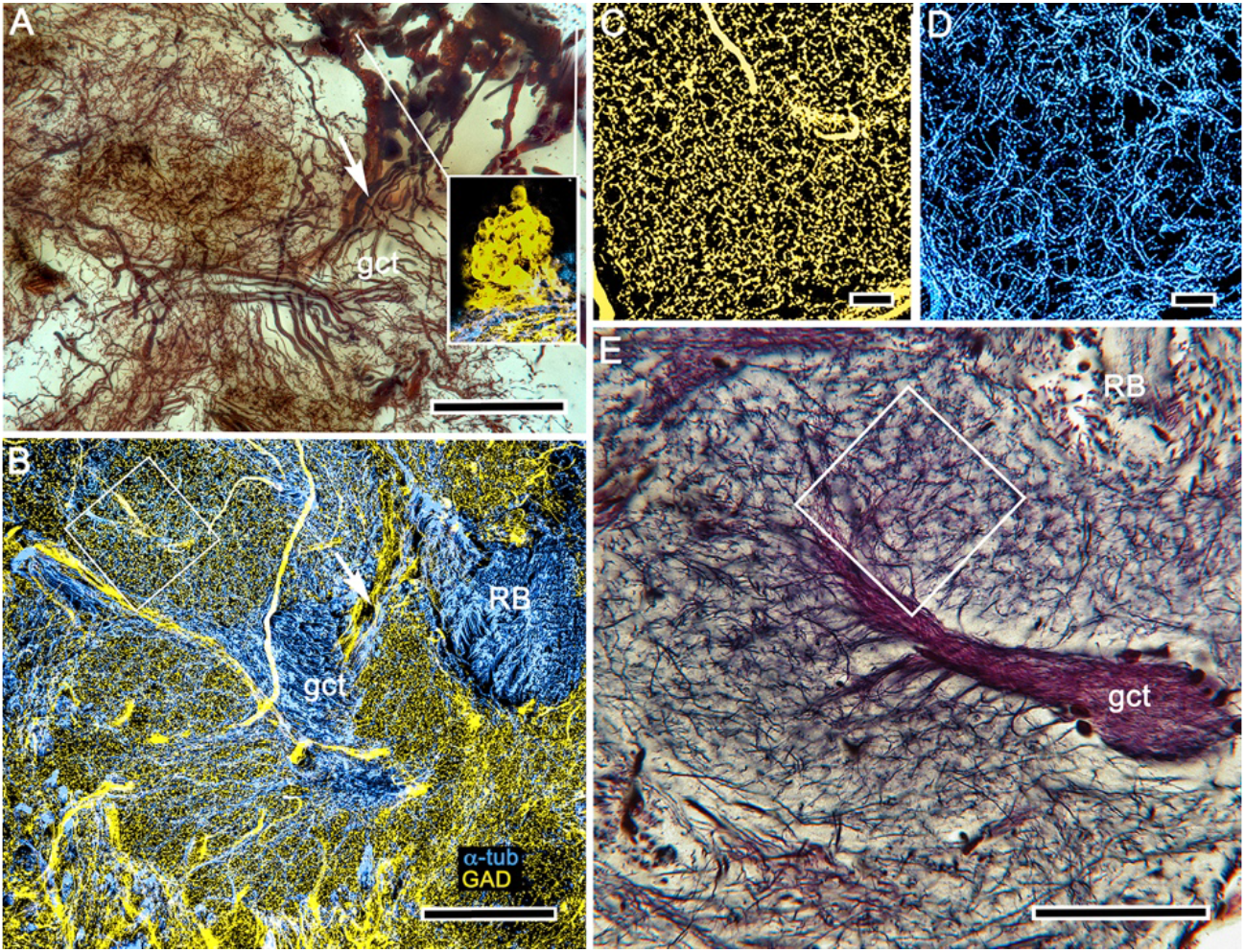
Evidence for local inhibition in the calyx by anaxonal neurons. (**A**) 40-50 large perikarya situated rostrally above the gyri are immunoreactive to glutamic acid decarboxylase (GAD, as shown in the right inset. These neuron cell bodies, which are here Golgi–impregnated, send stout cell body fibers (arrow) to the calyces where they provide widely branching and extremely dense processes that do not follow the patterns of the microglomerular mosaic. (**B**) GAD immunoreactivity reveals their branches and finest processes spreading through every calycal domain and hence positioned to potentially interact with all the intrinsic neuron dendritic fields. (**C, D**) GAD immunoreactivity in an area shown by the box in panel B, and an equivalent area in panel E showing the mosaic of microglomeruli. This mosaic is also evident from the pattern a α-tubulin-immunoreactive intrinsic neurons shown in D (same boxed area in B Scale bars, A, B, 100μm, C, D, 10μm, E, 50μm.

Whereas GAD immunohistology does not distinguish local calycal domains, these are indicated by the inhomogeneity of fields of 5HT- and TH-immunoreactive across the calyces (*Figure 6*). The neurons providing these fields also appear to be specific to the calyces and not part of a wider system that extends to other regions of the LPR. Those more distributed systems include densely branched arborizations that populate the optic lobe’s lobula and 5HT-immunoreactive neurons that arborize in certain neuropils of the CLPR as well as gyri. These are likewise distinct from the 5HT and TH-immunoreactive fields that intersect the mushroom body columns, which are described next.

**Figure 6.**
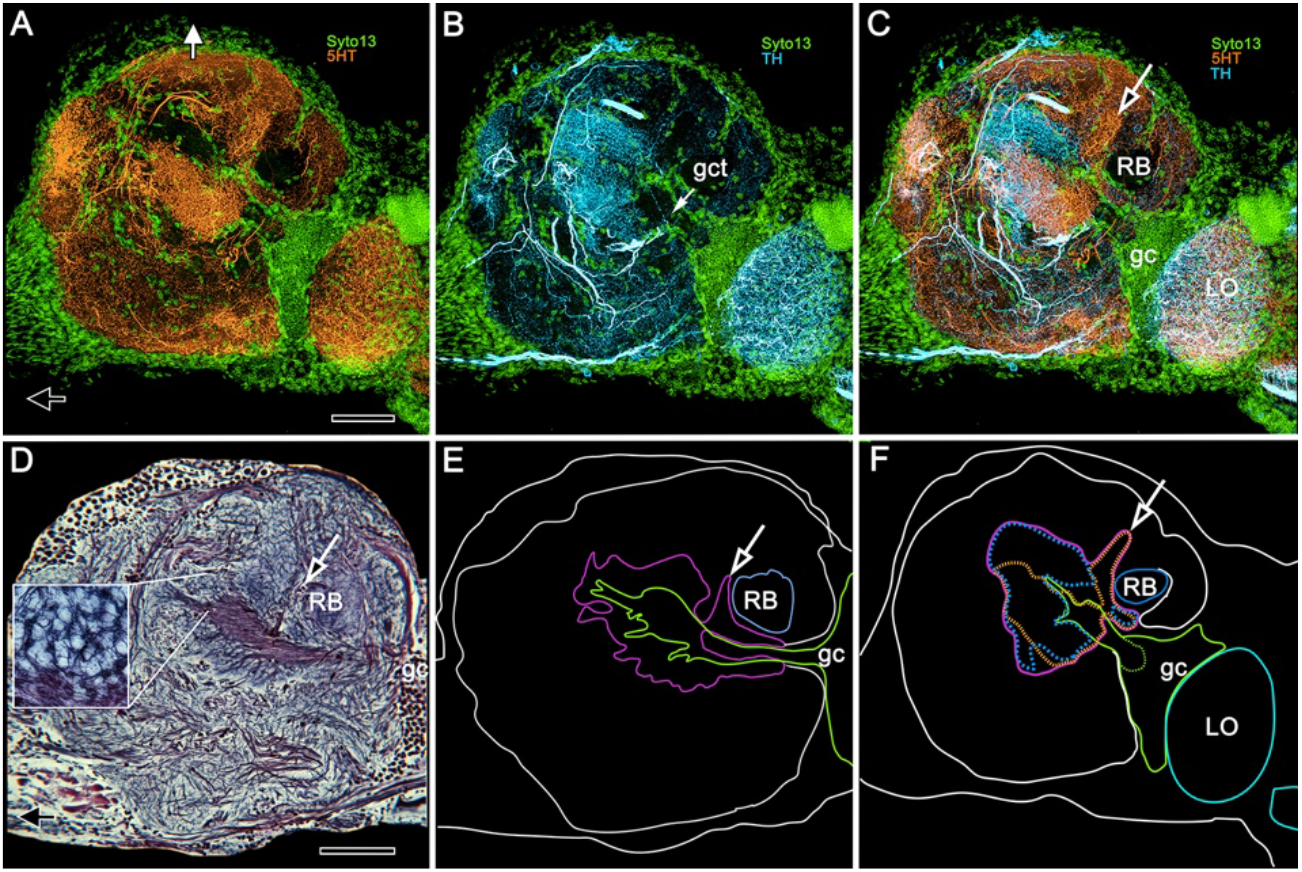
5HT and TH immunoreactivity define different calycal domains. (**A,B**) Although many volumes of the lateral protocerebrum are distinguished by 5HT (panel **A**, 5 hydroxytryptophan, orange) immunoreactivity, the densest system of branches is in a rostral domain of calyx 1. TH (panel **B**, tyrosine hydroxylase, cyan) defines two domains, one partially overlapping 5HT, the other not (panel **C**). (**D**) Bodian silver-stained calyx 1 showing its distinctive microglomerular mosaic (inset). (**E**) Outlined in purple is the total calycal perimeter at this level of sectioning and orientation. The open arrow indicates the origin of column 1. (**F**) Perimeter (purple) of calyx 1 in a section of the RLPR slightly rotated around its lateral-medial axis, mapping within it 5HT- and TH-immunoreactivity (as in panels A–C) also indicating the origin of column 1 (open arrow, as in panel C). Abbreviations: RB, reniform body; gc, globuli cell cluster; gct, globuli cell tract; LO, lobula. Scale bars for all panels, 100μm.

### 5. Transition from the calyces to the columns

Thus far we have demonstrated the composition of the calyces by approximately orthogonal networks provided by intrinsic neurons and their arrangements with microglomeruli.

Stomatopod and shrimp mushroom bodies demonstrate a clear transition of neuronal organization between the calyx and its columnar extensions (Wolff et al., 2017; Sayre and Strausfeld, 2018). In *H. nudus* there is no abrupt transition (*Figure 2)*. This ambiguity is seconded in brains treated with osmium-ethyl gallate, which show lateral protocerebral neuropil supplied by globuli cell neurites gradually fading from black to gray, signifying decreasing densities of neuronal membrane, finally giving way to the uniformly pale aspect of the overlying gyri (*Figure 1A, Figure2D,E*). 3-D reconstructions of the dark volumes show a layered arrangement at deeper layers of the calyces changing to a more corrugated appearance, particularly at its outer surface (*Figure 2, supplement 2*). Silver impregnations show that the base and some of the initial length of a column are clearly reticulated (*Figure 7C*). Double Golgi impregnation shows that these networks receive contributions from two sources: the extensions of axon-like processes from intrinsic neuron fields in the calyx; and intrinsic arborizations, many extremely complicated, that originate from thin neurites, presumed to belong to globuli cells (*Figure 7 - figure supplement 1*). An additional system of networks extends outwards to flank each calyx’s column (*Figure 7B*). These reticulations originate from a tangle of processes situated at a deeper level of the calyces (*Figure 7B*) that does not reflect any relationships with microglomeruli. The likely origin of this system (*Figure 7B*) is not from globuli cells but from a small clusters of cell bodies, some DC0-immunoreactive, situated ventrally close the inner margin of the lobula (*Figure 1C*). We interpret this tangle of processes (*Figure 7B*) as belonging to interstitial local interneurons providing elements to the columns at a level limited to their emergence from the calyces a small part of their initial length. The topology of these arrangements, including the small column-specific intrinsic arborizations (*Figure 7 - figure supplement 1E-K*) is reminiscent of interstitial neurons interposed between the calyces and pedunculus of the honeybee (Strausfeld, 2002).

**Figure 7.**
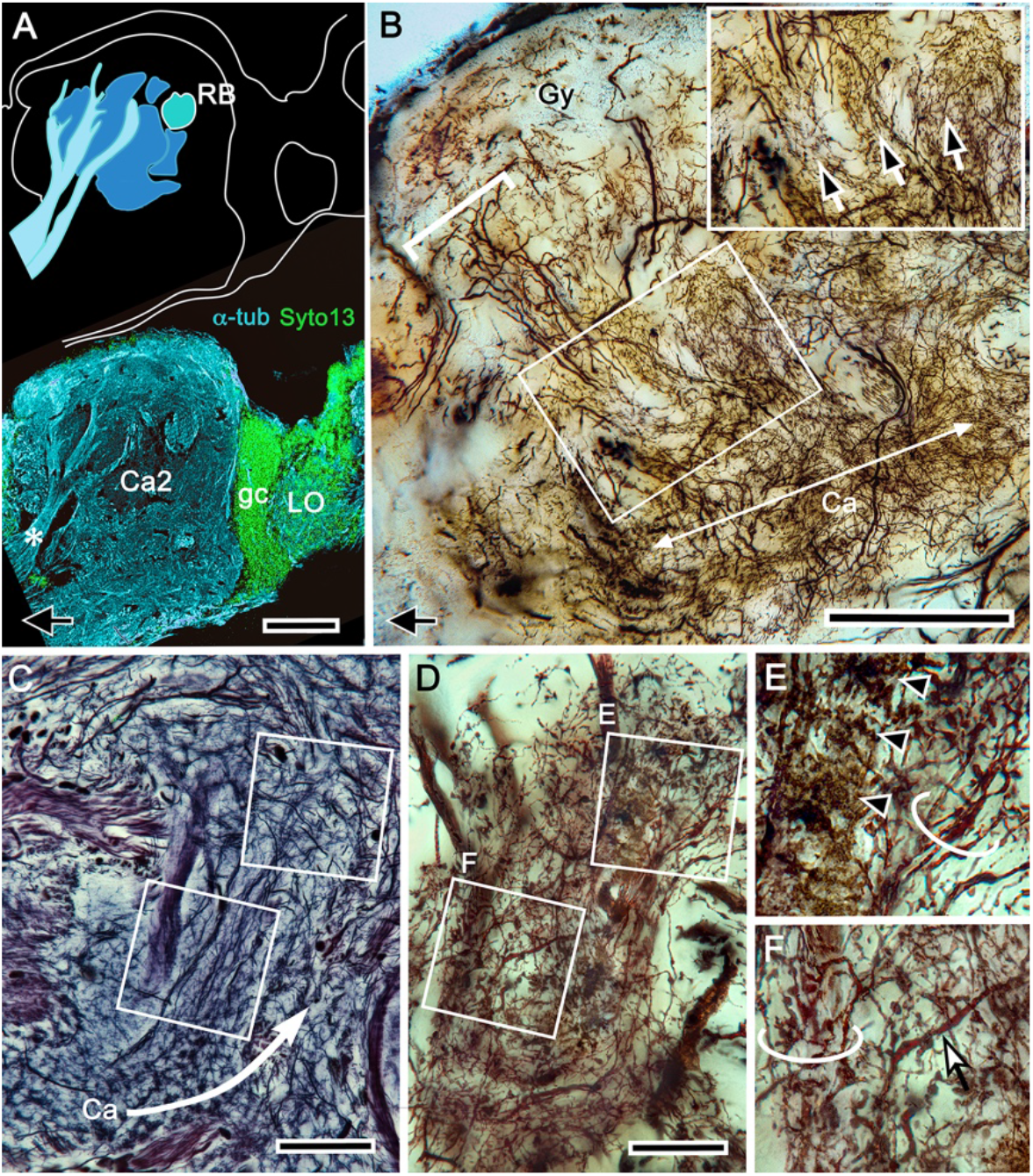
Origin of calycal extensions and columns. (**A, B**) Anti-α-tubulin (cyan) reveals the organization of large fiber bundles and coarse neuropil but, at low magnification, shows neuropils of the calyces as almost featureless (panel A). Part of the calyces are penetrated by incoming tributaries from the eyestalk nerve (asterisk in A, lower panel). A corresponding Golgi-impregnated region (B) demonstrates a system of medium-diameter branched processes that provide fine networks enwrapping the origin of columns. Some networks also extend as finger-like extensions outwards from the calyces. Panel B shows one column (upper left bracket) the base of which is flanked by those arrangements (enlarged in upper right inset indicated by arrows). This extensive system of processes does not derive from the globuli cell clusters although it is clearly associated with the entire extent of the calyces, suggesting a possible role in calyx-wide modulation. (**C-D**) As shown by this reduced silver stained section, the column arising from calyx 1 is extremely elaborate, comprising at least two longitudinal divisions each defined by arrangements of intrinsic neuron processes and their branched specializations. The curved arrow in panel C indicates the continuation of intrinsic neuron microglomeruli for a short distance alongside the column. The boxed areas in panel C, indicate corresponding levels identified in a Golgi-impregnated column 1 (D) showing enlargements in panels E, F, of their constituent arrangements. In panel E ascending fibers provide short collaterals (arrowheads) across one longitudinal subdivision of the column. In another subdivision, ascending intrinsic processes are arranged as a tangle (circled in panels E, F). Panel F also shows a large beaded branch (arrowed) of a mushroom body input neuron (MBIN) extending into the column. Abbreviations: RB, reniform body; Ca2, calyx 2; gc, globuli cells; LO, lobula; Gy, gyri. Scale bars, A, B, 100μm; C, D, 50μm. **Figure 7 ––– figure supplement 1.** Intrinsic neuron organization in columns **Figure 7 ––– figure supplement 2.** The interface between columns and gyri.

The origin of the columns and smaller extensions from the calyces are further demonstrated by selected levels of silver-stained brains. Serial sections resolve the uneven rostral border of the calyces and the two columns reaching outwards towards the overlying gyri (*Figure 7 - figure supplement 2*). The series also demonstrates the fidelity of silver and osmium-ethyl gallate in showing corresponding origins of the columns from the two calyces. These aspects emphasize that the seemingly irregular morphology of the rostral part of the calyces is not irregular by happenstance but is a defining feature of the lateral protocerebrum.

Reduced silver preparations demonstrate that a column is subdivided into parallel components as in insect mushroom bodies, where each subdivision has a distinctive arrangement of intrinsic neurons (Strausfeld et al. 2005; Sjöholm et al., 2005; Tanaka et al., 2008) and is defined by distinctive patterns of gene expression (Yang et al., 1995). What is unexpected is the degree of elaboration observed in the varunid mushroom body. Golgi impregnations demonstrate that, compared with other pancrustaceans, mushroom bodies in a crab are stunningly complex, both with regard to their calyces and columns. The varunid’s columns consist of parallel intrinsic cell fibers that are almost obscured by numerous other arrangements, the variety of which is compatible with the variety of their morphological fields in the calyces. Comparison of the silver-stained column originating from calyx 2 to one impregnated by mass Golgi impregnations (*Figure 7C,D*) demonstrates a discrete and complicated system of processes, some providing bridgelike connections across them (*Figure 7E*), others having irregular branches, lateral prolongations, and bundled parallel fibers (*Figure 7E,F*). In addition, larger dendrite-like elements arising from stout processes within the columns suggest that those processes may provide outputs extending outwards to the overlying gyri (*Figure 7D,F*), a feature to which we return in section 7.

Golgi impregnation has its own peculiar limitations: it is stochastic, revealing probably less than 1% of the total population of neurons, even after double impregnation. Nevertheless, mass impregnation further suggests that the columns from the two calyces are not identical. Although networks of intrinsic cells in both columns receive terminals from branches of the olfactory glomerular tract, there are clear differences between the columns from calyx 1 and calyx 2. Column 2 is broader and more difficult to discern in reduced silver. In part because many of its intrinsic processes are all extremely thin, many less than a micron in diameter (see section 7 below). At first sight, the organization of processes comprising column 2 appears to be chaotic. However, when scrutinized more closely the organizations of its dendrites and collaterals contribute to subsets of different arrangements disposed across the width of a column and at different levels along it (*Figure 7 supplement 1A-C*). Reciprocal channels to the column from the gyri are suggested by the presence of descending axons flanking the column, providing varicose collaterals into it (*Figure 7 - figure supplement 1A*). These varieties of columnar arrangements are further considered in the Discussion.

Fields of 5HT- and TH-immunoreactive dendrites extend across columns. Here we show fields relating to column 1 from calyx 1 that provide clear resolution of their arrangements from the calycal origin to the interface with gyri (*Figure 8*). Fields are arranged at specific levels along the column and they intersect different domains across the column, as occurs across the parallel subdivision of the columns (lobes) of mushroom bodies in *Drosophila* and other insects (Nässel and Elekes 1992; Liu et al., 2012; Aso et al., 2014a; Hamanaka, 2016; Tedjakumala et al., 2017). The first of these fields occurs at the origin of the column from the calyx (*Figure 8A,B*) above where the neat orthogonal arrangements seen at deeper levels give way to the first appearance of looser arrangements, including parallel fibers (*Figure 8C,F*). Anti-5HT- and anti-TH-immunoreactive fields are clearly segregated (*Figure 8A and D*) at or near the column origin but begin to overlap at more rostral levels closer to the gyri, at locations where osmium-ethyl gallate preparations show the columns branching, with tributaries extending to overlying gyri. At those levels, immunoreactive processes appear to spread laterally into the gyri themselves (*Figure 8G,H*).

**Figure 8.**
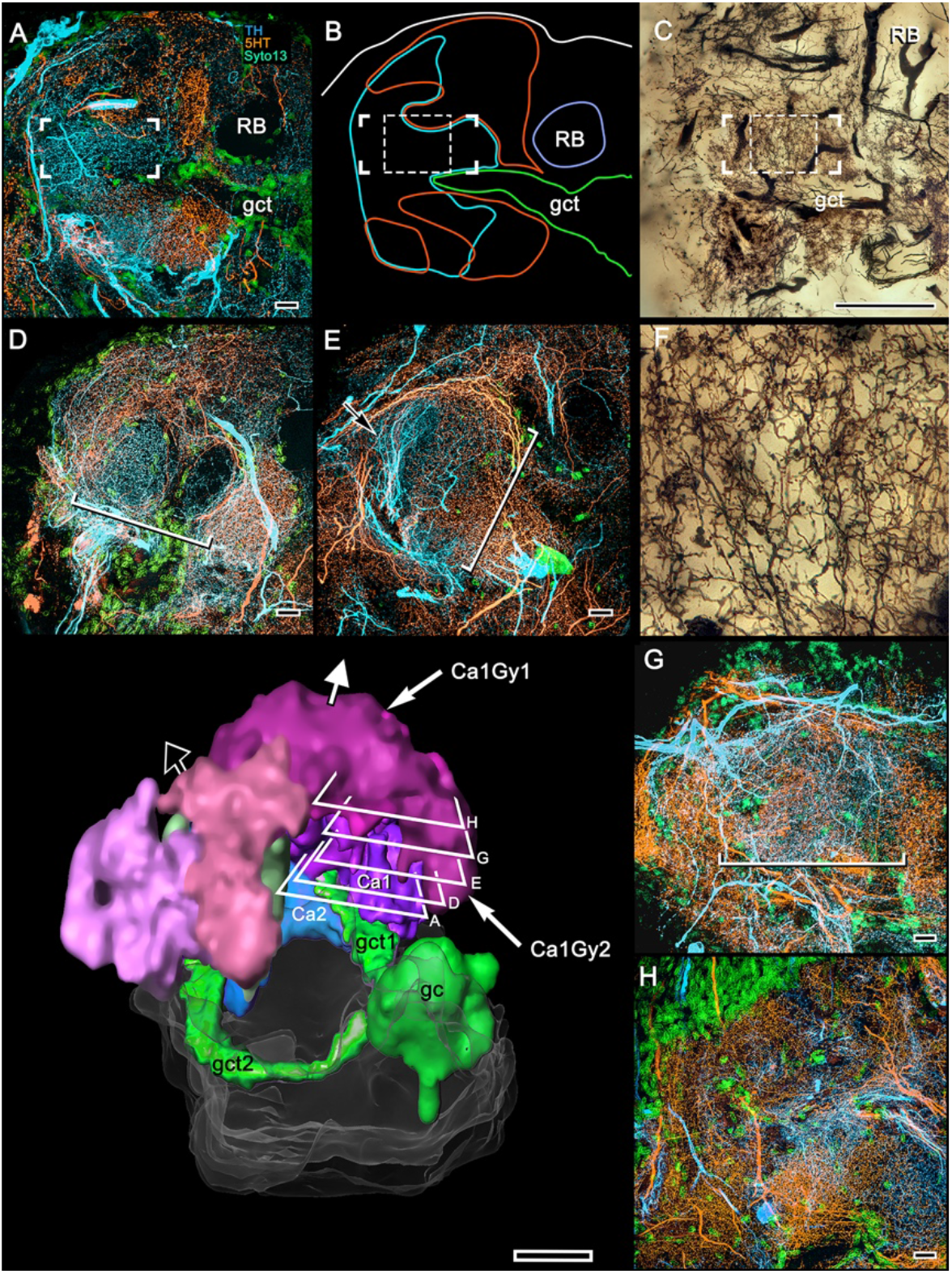
5HT and TH immunoreactivity resolve different levels in the calyx 1 column. Looking downwards through the rostral surface of LPR, this sequence of confocal slices (**A,D,E,G,H**) shows views starting at the column’s origin from calyx 1 (panel **A**) then following its outward passage rostrally to the level at which the column merges with the overlying gyri (panels **G**, **H**). (**B)** shows outlines of 5HT- and TH-immunoreactive fields at level A in relation to the globuli cell tract (gct, green) and the cross section of the reniform body’s pedestal (RB).The dendritic field of a TH-immunoreactive cell is bracketed in A,B and a smaller portion of it is denoted by a dashed square in B. (**C**) Golgi preparation of dendrites in area equivalent to bracketed and dashed areas in (B) show that at this level, networks of intrinsic neurons have given way to more loosely defined reticulations. (**F**) Enlargement of dashed square in (C). At the level shown in panel A, 5HT- and TH-immunoreactive fields are segregated across the origin of the column and continue to be, at least until the level shown in panel **E**, suggesting 5HT- and TH-immunoreactive fields are restricted to the column’s longitudinal subdivisions. TH-immunoreactive fields can be noticeably clustered within a small area of the column’s cross-section (arrow in E) as are MBONs in *Drosophila* mushroom bodies. The width of the column (brackets in panels D, E, G) increases gradually through layers **D**, **E**, and **G**(see brackets), and in the gyrus (**H**) its borders are no longer distinguishable. The 3–D reconstruction (lower left) shows a view looking into the lateral protocerebrum from its confluence with the optic lobes, its medial axis indicated by the black arrow and the rostral axis by the white arrow pointing upwards. Confocal levels are indicated as well as the two gyri (Ca1Gy1, Ca1Gy2) associated with the Ca1 column and the globuli cell tracts to Ca1 and Ca2 from the globuli cell cluster (gc). Other abbreviations: gct, globuli cell tract; pds, pedestal of the reniform body; Gy, gyrus. Scale bars, A-E, G, H, 10μm; C and 3–D reconstruction, 100μm.

### 6. Organization of gyriform neuropils

Five distinct gyri comprise the outer layer of the RLPR (*Figure 9A*). But only four gyri are associated with the two calyces. A fifth proximal gyrus situated near the entry of the eyestalk nerve is notable for its neuronal organization distinguishing it as entirely separate from the mushroom body (*Figure 9A,F*).

**Figure 9.**
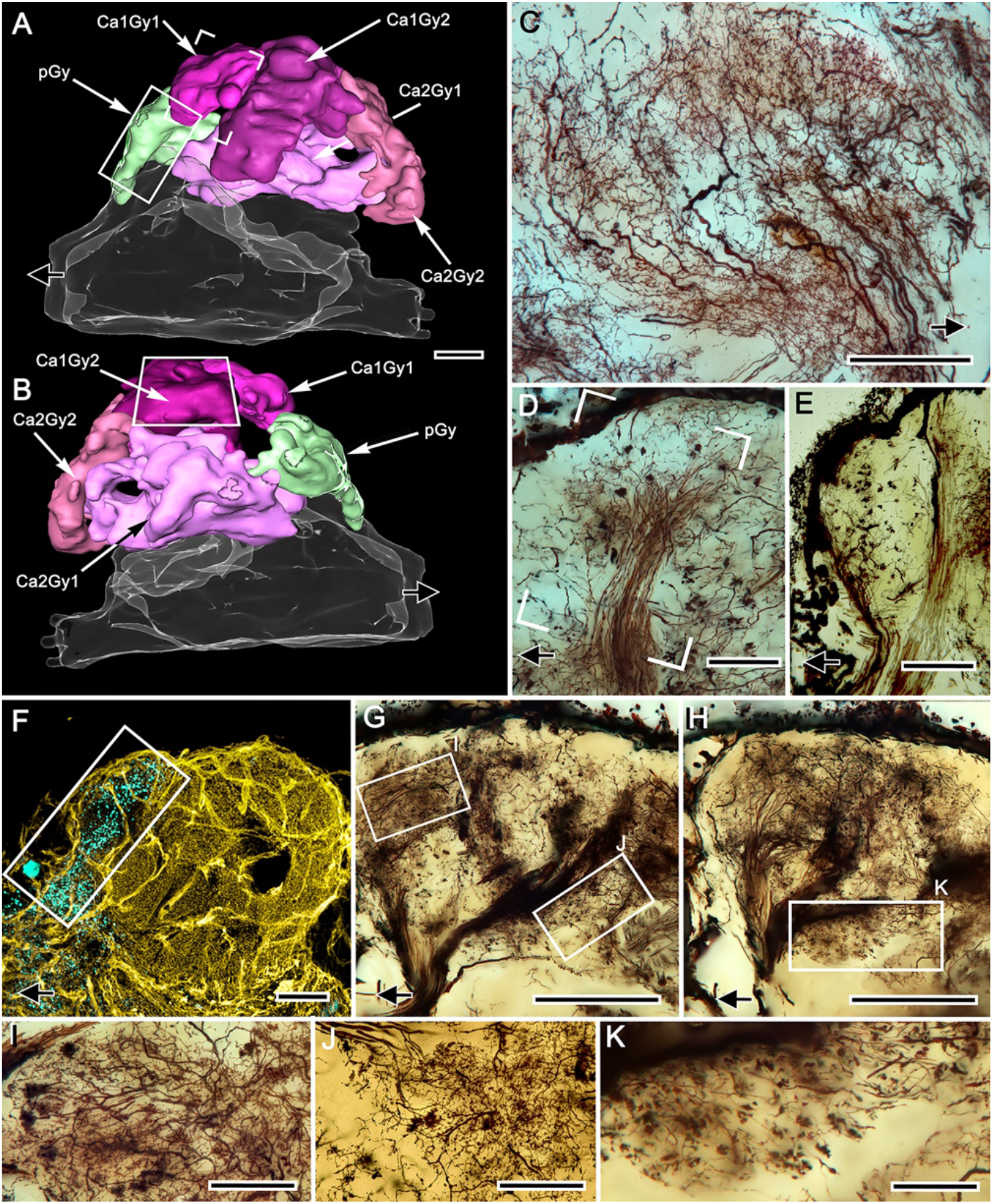
Gyriform neuropils: afferent and efferent organization. (**A, B**). Amira-generated 3–D reconstructions of the gyri and their calyces. Black arrows indicate the direction of the eyestalk nerve; likewise in panels C–H. (**C**) Ensemble of efferent dendrites in gyrus 2 associated with calyx 1 (Ca1Gy2). The represented area is boxed in panel B. (**D**) Afferent tributary from the antennoglomerular tract (olfactory projection neurons) suppling gyrus Ca1Gy1 neuropil (open rectangle in panel A). (**E**) Smooth oval presynaptic specializations denote the terminals of another antennoglomerular tributary that exclusively supplies the most proximal gyrus (pGY, rectangle in panel A), which has no obvious connection to the mushroom body. (**F**) Double-labelling with anti–GAD (yellow) and anti–allatostatin (cyan) of the lateral protocerebrum resolves allatostatin immunoreactivity in the proximal gyrus (rectangle in A and F), which shows little to no GAD immunoreactivity. (**G, H**) Stacked z-axis projections of Golgi-impregnated tracts, imaged through two successive 50-μm sections showing massive supply by afferent neurons. These reach levels of the lateral protocerebrum that include those at and well beneath the gyriform layer (box I, panel G), including levels of the calyces (box J in panel G and box K in panel H). Notably, many efferent neurons relaying information from gyri (panel H) send their downstream axons from the lateral protocerebrum into these tracts. (**I-K**) Enlargements from regions indicated in G, H, but at sample thicknesses of less than 20 μm to isolate details obscured in flattened optical sections. Scale bars, A–C, F–H, 100μm; D,E, 50μm; I– K, 50μm. **Figure 9 ––– figure supplement 1.** Perikaryal distributions and gyri neurons

Mass impregnations demonstrate at this outer level of the lateral protocerebrum large numbers of efferent neurons and their dendritic processes, many of which originate from clusters of loosely arranged patches of perikarya that extend over the outer surface of the lateral protocerebrum (*Figure 9 – figure supplement 1*). Although stochastically impregnated, the abundance and variety of these neurons suggests the gyri contain dense arrangements of efferent (output) and local interneurons (*Figure 9-figure supplement 1D-I*). Golgi impregnations reveal swathes of axons extending across the gyri, suggesting lateral interactions amongst them. Oblique palisades of dendritic trees, which have branching patterns peculiarly reminiscent of vertebrate pyramidal cells (*Figure 9B*), send axons to fascicles that extend to the midbrain via the eyestalk nerve.

Like deeper levels of the lateral protocerebrum, gyri comprising this superficial level of the lateral protocecerum are supplied by terminals of afferent neurons, the axons of which radiate out from the eyestalk nerve. Many of these terminals are identified as originating from discrete fascicles that comprise the antennal globular tract, denoting not only their identity as axons of projection neurons from the olfactory lobes but suggesting that different classes of projection neurons comprise different fascicles (*Figure 9D,E*). The terminals of projection neurons are denoted by their swollen boutons, the different morphologies of which are distinctive. For example, terminals spreading out over the surface of gyrus 1, belonging to calyx 1, have rough ball-shaped profiles that are distinct from larger smooth boutons from extremely slender axons supplying the fifth proximal gyrus (pGy in *Figure 9A,B,E*). That gyrus’s large allatostatin-immunoreactive processes and the paucity, even absence, of GAD-immunoreactive networks suggests its complete independence from the mushroom body’s organization (*Figure 9F*). Nor, as shown by osmium-ethyl gallate, is this gyrus associated with any of the calyces’ outward extensions.

In addition to axons extending from the eyestalk nerve providing afferents to superficial levels in the gyri, all levels in the rostral protocerebrum receive an abundant supply of terminals, including many penetrating to deep levels of the calyces (*Figure 9G-K*). Again, it is worth bearing in mind when viewing the images in Figure 9G, H that Golgi impregnations likely reveal not more than a few percent of the constituent neurons providing these already dense arrangements of incoming fiber tracts.

### 7. Transitions to gyri

We have shown that four gyri overlie two large columns, each extending from one of the two calyces, as do occasional smaller finger-like extensions. Osmium-ethyl gallate demonstrates that the columns bifurcate close to the gyri (*Figure 1A, 2D,E*), a feature also resolved in serial reduced-silver-stained sections (*Figure 7 - figure supplement 1A-H*). However, neither method allows the identification of a clear inner margin of a gyrus or an obvious termination of the column. Here we consider neural arrangements that denote the ambiguity of the interface between the mushroom body’s columns and their corresponding gyri.

Tangential sections across the gyri demonstrate novel and as yet puzzling aspects of their composition (*Figure 10A,B*). One is the presence of dendritic trees, as in Figure 10B, here represented by a single impregnated neuron, the axon of which groups with others extending to the lateral protocerebral root of the eyestalk nerve. The dendrites are seen overlapping with elaborate networks of finely branching processes that terminate in small bead-like swellings suggestive of presynaptic specializations. This network (*Figure 10B*) derives from extremely thin processes that ascend into the gyrus from outward extensions of the calyx (bracketed in *Figure 10B*). Further examples (*Figure 10C*) show that the dendrites of a gyrus’s efferent neurons overlap other arrangements of putative ascending presynaptic fields. In this example, the fields do not appear to be part of an intervening synaptic network but instead their disposition with the efferent dendritic trees suggests that they likely terminate there (*Figure 9C*). In addition, highly restricted systems of mixed spiny and beaded arborizations suggest local anaxonal neurons that may provide synaptic interfaces between such terminal systems arising from the columns and the dendrites of gyral efferent neurons (*Figure 10D,E*). Evidence that arborizations predominantly equipped with presynaptic specializations likely originate from the calyces is indicated by densely packed elements situated distally in the columns. These are equipped with short digiform specializations arranged as small basket-like receptacles that clasp afferent terminals of unknown origin (*Figure 10F-H*).

**Figure 10.**
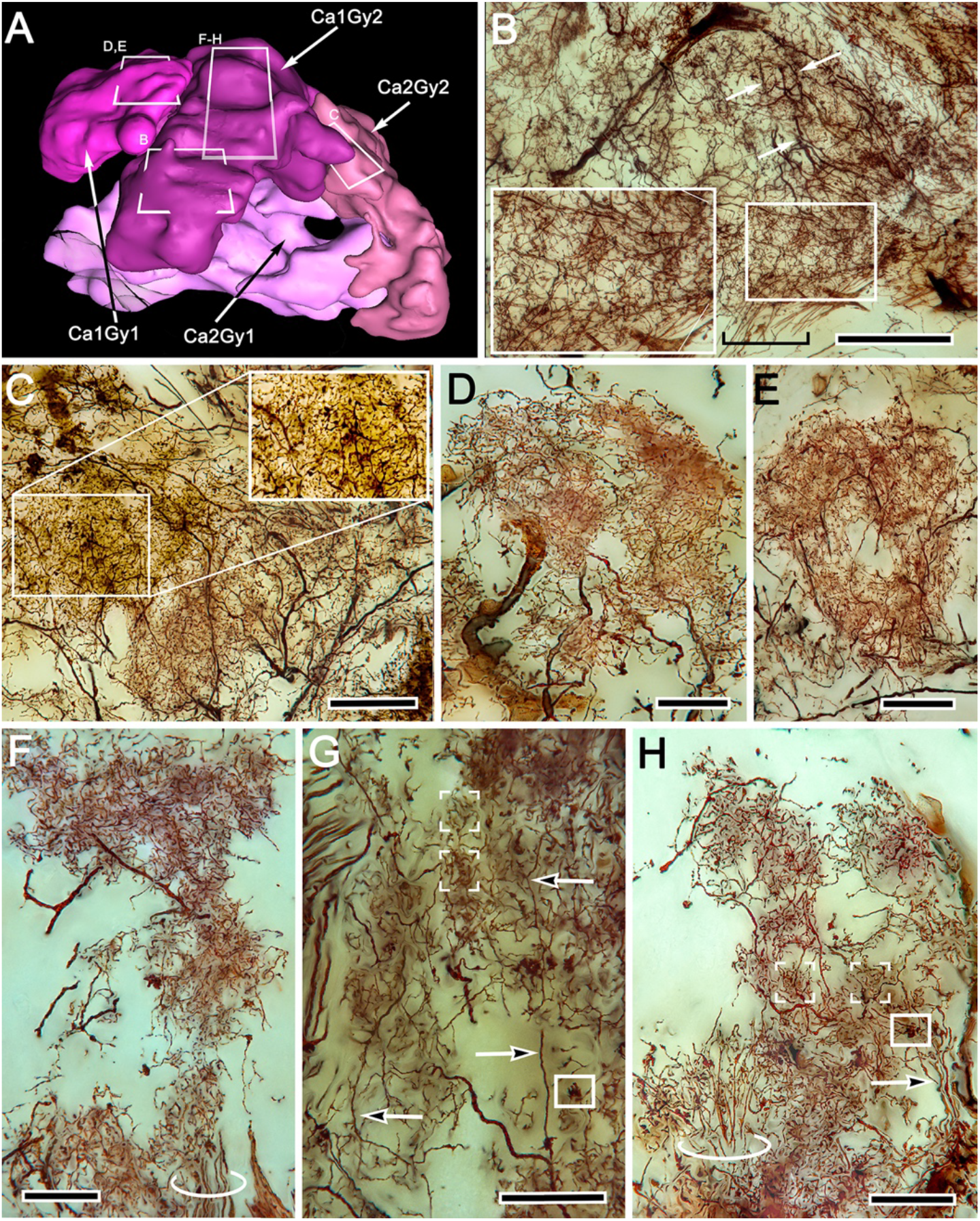
Gyrus neuropils: local circuit neurons. (**A**) 3-D reconstruction of gyri associated with calyx 1 and 2, showing approximate locations of networks shown in B–H. (**B**) Dense network supplied by ascending processes (boxed area is enlarged on left), ascribed to Ca1Gy2 derived from column 1. Dendrites of an efferent neuron (arrows) spread through this layer. (**C**) Efferent dendrites overlap a broad field of dense terminal specializations. Gnarled bouton-like specializations (boxed and enlarged on right) show that terminals from the eyestalk nerve enter this level. (**D-H**) Dense and elaborately branched elements interpreted as local interneurons (D, E) and intrinsic elements of the column (F-H). Ascending processes (circled in F,H and arrowed in G,H) are further evidence of local circuits. Dense groups of converging processes (box brackets in G,H) correspond in size to afferent boutons (boxed in G,H). Scale bars, B, C, 100μm: D-H, 50μm. **Figure 10 ––– figure supplement 1.** Column-gyrus interface organization.

The rare Golgi impregnation that resolves one of the columns as a recognizable ensemble of heterogeneous elements within the column’s boundaries also demonstrates that its terminus is “open,” meaning contiguous with the gyral neuropil into which some of its fibers project (*Figure 7-figure supplement 1A; Figure 10 – figure supplement 1A*). Also fortuitous are preparations that demonstrate the terminus of a column coinciding with bundled endings from the antennoglomerular tract. Such a preparation (*Figure 10 supplement 1B*), also shows that the terminus of the column is open to gyral neuropil around it and that processes extend from the column into the gyrus. Combined labelling with anti-synapsin and actin at the terminus of the column (*Figure 10 - figure supplement 1C*) further indicates that sizes and densities of sites indicating synaptic convergence differ systematically across it. The flanking gyral neuropils also betray substantial variation in synaptic densities. Inputs to the gyri do not, however, derive solely from calycal columns and afferents from the antennoglomerular tract. Reduced silver shows that tracts from caudal neuropil associated with the optic lobe extend rostrally to the superficial levels of a gyrus (*Figure 10 - figure supplement 1D*), yet another indication of the complexity of organization and its multimodal nature even at the most distal level of the mushroom body. The possible significance of this is considered below in the Discussion.

### 8. The reniform body

We have shown that the lateral protocerebrum is divided into a rostral and caudal part (RLPR and CLPR), the former containing the paired mushroom bodies. The mushroom bodies are not, however, the sole components of the RLPR. The pGy gyrus nearest to the RLPR’s junction with the eyestalk nerve is distinct and probably to some degree functionally separate from gyri associated with the mushroom bodies (*Figure 9F*). In addition, there is another prominent system of circumscribed neuropils situated close to the lateral border of the RLPR, arising from a dorsal group of relatively small perikarya almost adjacent to the optic lobes. These neuropils comprise the reniform body, a center identified in stomatopods by Bellonci (Bellonci, 1882), who distinguished it from the very prominent mushroom bodies in that taxon. The same center has been described in detail in the stomatopods *Neogonodactylus oerstedii*, *Gonodactylus smithii*, and *G. chiragra*, and the carideans *Lebbeus groenlandicus* and *Alpheus bellulus.* In all these species the reniform body adopts the same morphology: a densely populated bundle of axons, referred to as the pedestal extends from the dorsal surface of the RLPR to its ventral surface, providing four discrete neuropils called the initial (dorsal), lateral, and the ventrally situated proximal and distal zones (Wolff et al., 2017; Thoen et al., 2019; Sayre and Strausfeld, 2019; Strausfeld et al., 2020). In all of the cited taxa, the reniform body is situated between the columnar mushroom body and the lobula.

Precisely the same placement is observed in the varunid lateral protocerebrum (*Figure 11*). The reniform body occupies a location lateral to the ensembles of neurons that constitute the mushroom body gyri (*Figures 1,11*). Reconstructed from serial osmium-ethyl gallate sections, the reniform body’s pedestal lies immediately above, i.e., rostral to, the two tracts of neurites from the globuli cell cluster (the globuli cell tract) that provide intrinsic neurons to the mushroom body calyces (*Figure 11 C*). In *H. nudus*, the reniform body pedestal extends from a mass of cell bodies at the dorsal surface and extends almost to the lateral protocerebrum’s ventral surface, bifurcating into two main tributaries about half way along its length (Figure 11 F). These give rise to the lateral, distal and proximal zones, each zone composed of substantial volumes of arborizations (*Figure 11 D*). The reniform body is thus entirely distinct from the mushroom bodies, and is situated distally with respect to their centrally disposed calyces (*Figure 11 E*). The reniform body’s thick pedestal, albeit columnar in form, is distinct from a mushroom body column in that its constituent axons are naked; lacking spines or other specializations that define a mushroom body. The reniform body’s four zones of branching collaterals contribute to structures reminiscent of microglomeruli. However, they are of larger girth and synapsin-actin labelling shows them to have significantly fewer converging presynaptic elements than a calycal microglomerulus (*Figure 10, figure supplement 1D,E*). Despite these clear differences the reniform body is strongly immunoreactive to anti-DC0 showing that it also is a learning and memory center as demonstrated by Maza et al. (*Maza et al., 2016*).

**Figure 11.**
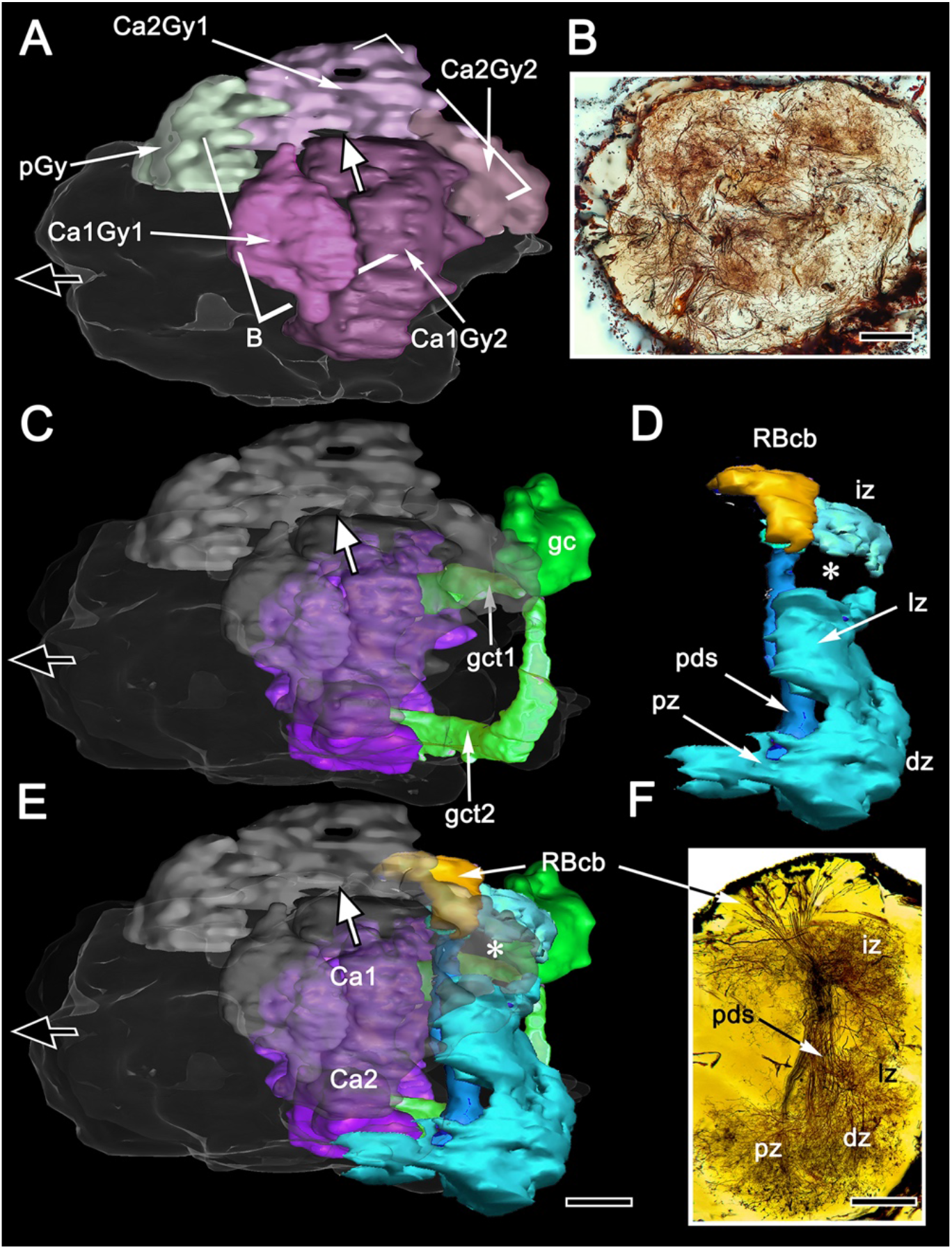
Disposition of the reniform body. (**A**) 3-D reconstruction of gyri from osmium-ethyl gallate specimens identifies their 2:1 relationship with columns and their total volume, which can be compared to the distribution of gyrus neurons resolved by Golgi impregnations (**B,** level indicated in A). The Golgi method impregnates a fraction of the constituent neurons, the fields of which are mostly constrained to a gyrus. Here impregnated neurons are viewed through the LPR’s rostral surface. In panel A (and panels C E) the black arrow indicates the medial axis, the white arrow the rostral axis. (**C**) The globuli cell cluster (gc) provides two globuli cell tracts (gct1, gct2) to the two mushroom body calyces Ca1, Ca2. The lateral lobe of gyrus 2 associated with calyx 2 (indicated by the asterisk in panel E) reaches out distally and fills a gap (asterisk in D) between two processes (the intermediate and lateral zones, iz and lz) originating from the reniform body’s pedestal (pds). (**D**) The reniform body pedestal is a dense fiber bundle originating from a dorsal cluster of small cell bodies (RBcb) on the distal flank of the RLPR. The pedestal’s ventrally dividing branches, shown Golgi-impregnated in panel **F**, provide the dorsal and proximal zones (dz, pz). (**E)** Combined volumes in A,C, and D demonstrating the position and modest dimensions of the reniform body relative to the mushroom bodies, comprising the calyces (Ca1, Ca2) and their columns and associated gyri. Scale bars, A-C, B, F, 100μm. **Figure 11 ––– figure supplement 1**. Comparison of synaptic densities associated with the calyces and reniform body.

## Discussion

### Unique organization of the varunid mushroom body

Studies by Hanström in the 1920s and 1930s on the brains of malacostracan crustaceans concluded that mushroom bodies equipped with calyces and columns (lobes), as they are in insects and mantis shrimps (Bellonci, 1882), have been reduced in decapods to columnless centers lying immediately beneath the rostral surface of the lateral protocerebrum. Recent comparisons of the brains of reptantian decapods have confirmed that its lineages share the evolutionary trend of reduction and loss of the mushroom body column (Strausfeld and Sayre, 2019). Those mushroom bodies nevertheless retain most elements of ancestral networks: diffusely arranged in Axiidea, Astacidea, Achelata (Strausfeld and Sayre, 2019), but precisely stratified in terrestrial and marine Anomura, (hermit crabs: Harzsch and Hansson, 2008; Wolff et al., 2012; Strausfeld and Sayre, 2019).

Ocypodidae (fiddler crabs) and Varunidae (shore crabs), two lineages of Brachyura (true crabs) do not conform to this trend. As demonstrated previously and again here, large DC0-positive domains cover much of the varunid lateral protocerebrum, suggesting relatively enormous (for an arthropod) learning and memory neuropils (Strausfeld et al., 2020). The proposition that the reniform body is the crab’s mushroom body (Maza et al., 2016, 2020) is refuted by Golgi impregnations and 3-D reconstructions of the lateral protocerebrum demonstrating the reniform body as entirely distinct from the huge DC0-positive mushroom body adjacent to it (*Figure 11*).

Similarity, conjunction, and congruence are established criteria for assessing phenotypic homology (Patterson, 1988). In all the lineages where reniform bodies have been identified these criteria exclude it as mushroom body homologue. That the brachyuran mushroom body has until now remained unrecognized can be attributed to its hiding in plain sight as an unexpected and radical reorganization. Even its population of conservatively estimated 22,000 densely packed globuli cells is not where it would be expected to be. Based on observations of other pancrustaceans, globuli cells are clustered over the rostral surface of the lateral protocerebrum, close to where it join the eyestalk nerve (see Sayre and Strausfeld, 2019). Amplifying this distinction is that the entire mushroom body is inverted such that its two voluminous calyces are deeply buried in the rostral domain of the lateral protocerebrum where they receive inputs from the deutocerebrum’s olfactory lobes and the protocerebrum’s optic lobes.

We show here that each calyx gives rise to a large column extending outwards and merging with broad overlying gyri. The gyri are pillowed into folds and sulci thus forming a cortex-like architecture. This system receives additional afferent supply and gives rise to pyramidal-like ensembles of efferent neurons, the axons of which extend to the eyestalk nerve. Throughout this massive structure, the mushroom body and its gyri are strongly immunoreactive to antibodies raised against DC0.

Although the crab mushroom body appears vastly different from the right-way-up “traditional” insect or stomatopod homologue (Figure 12), it nevertheless shares with them their defining traits. For example, intrinsic neurons originating from globuli cells contribute to the brachyuran calyces and columns as do the Kenyon cells of an insect mushroom body. As in stomatopods, shrimps, and insects, the axon-like processes of intrinsic cells are clustered in their respective fascicles, as they are in any one of the longitudinal divisions of the insect mushroom body columns, where short intermingling collaterals can provide orthogonal connectivity (Strausfeld et al. 2005; Sjöholm et al., 2005; Tanaka et al., 2008). Other longitudinal divisions of the varunid mushroom body column are populated by intrinsic processes that interweave to provide rectilinearity, as they do in one of the two mushroom body columns of the shrimp *Lebbeus groenlandicus* (Sayre and Strausfeld, 2019), or in the stratified dome-like mushroom body of hermit crabs (Wolff et al., 2015). We show here that true crabs express the whole panoply of intrinsic cell arrangements. These include segregated and intermingling “parallel fibers” as in Stomatopoda or the core lobe (column) of *Drosophila* (Wolff et al., 2017; Strausfeld et al., 2003); rectilinear networks that extend part way into a column from the calyx; dense arrangements of intrinsic processes extending across the columns comparable to certain neurons populating the mushroom body column in honey bees or crickets (Strausfeld 2002; Hamanaka and Mizunami, 2019); and arrangements comparable to the longitudinal laminations of the mushroom body lobes (columns) of Dictyoptera and Lepidoptera (Sinakevitch et al., 2001; Sjöholm et al., 2005). In short, the crab mushroom body column appears to embody all known arrangements across Pancrustacea. If each arrangement of a specific type of intrinsic neurons represents a distinct computational property, as suggested from studies of honeybees and fruit flies (Traniello et al., 2019; Perisse et al., 2016), then the crab would appear to be singularly well-equipped. The same applies to its calycal arrangements, which in *H. nudus* are elaborate and discretely partitioned into territories denoted by different morphologies of intrinsic neurons and afferents. This zonal organization corresponds to the more compact and highly ordered modality-specific domains typifying hymenopteran calyces (Gronenberg 2001), and, to a lesser extent, *Drosophila* calyces as well (Yagi et al., 2016; Li et al., 2020).

**Figure 12.**
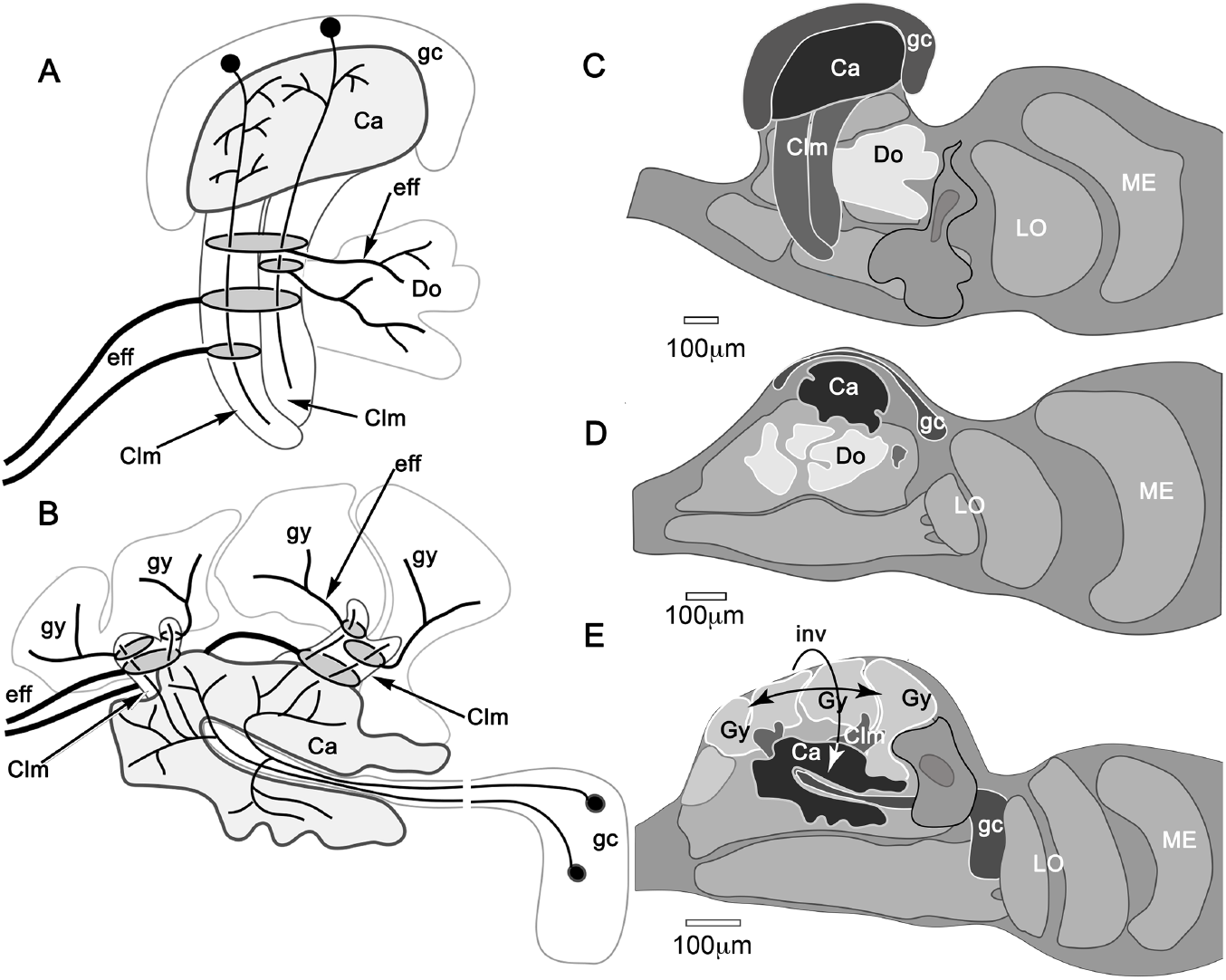
Mushroom body inversion and gyriform organization. (**a, b**) Schematics of corresponding neuron ground patterns. (**a**) In Stomatopoda, intrinsic neuron dendrites corresponding to the Kenyon cells of insect mushroom bodies occupy specific domains (here levels) in the calyces. Their prolongations down the columns are intersected by efferent dendritic fields, the axons of which extend to the midbrain (left) or to domains (Do) deeper in the lateral protocerebrum. (**b**) In *H. nudus*, the mushroom body is inverted. Intrinsic neuron dendrites occupy characteristic planar territories in the calyces, with rostrally projecting extensions in the columns intersected by efferent neuron dendrites. As in stomatopods (TH-MBONs), axons of some efferent neurons extend medially to the superior protocerebrum of the midbrain (not shown). Other efferents supply the gyriform neuropils overlying the lateral protocerebrum (Gy). (**c-e**) Comparisons of mushroom bodies in Malacostraca. (**c**) In the stomatopod *N. oerstedii* globuli cells cover the calyx; columns project caudally into the lateral protocerebrum neuropil. Neuropils within the lateral protocerebrum (Do) receive inputs from the lobes. (**d**) In the crayfish *Procambarus clarkia* the lobeless mushroom body lies at the lateral protocerebrum’s rostral surface. Neuropils in the lateral protocecerum receiving its outputs. (**e**) In *H. nudus*, calyces are deeply buried in the lateral protocerebrum. Columns extend rostrally to an overlying gyriform neuropil with the entire arrangement indicating mushroom body inversion and lateral expansion of the rostral gyri. Abbreviations: Ca, calyx; Clm, mushroom body column; Do, domains receiving mushroom body efferents; eff, efferent neurons; gc, globuli cells; Gy, gyri; GyC, gyriform cortex; LO, lobula; ME, medulla.

The varunid lateral protocerebrum has two calyces, each providing a massive column. This doubling of the calyx has many precedents. In the stomatopod lateral protocerebrum there are four adjacent columns, each associated with a distinct ensemble of globuli cells and hence a calyx. That organization has been compared to the *Drosophila* mushroom body, which is a composite neuropil of four hemi-mushroom bodies derived from four clonal lineages, as it is in the honey bee (Wolff et al., 2017; Ito et al., 1997; Farris et al., 2004). Consequently, the mushroom bodies in an insect’s lateral protocerebrum are paired but completely fused. The varunid has two calyces in immediate contact, but not quite fused as no intrinsic neuron extends its field of processes into both. Only one calyx (CA2) appears to receive input from a tract of axons carrying relays from the medulla and from the reniform body. Such an asymmetric input to twin calyces is not unique: a comparable arrangement is found in the *Drosophila* mushroom body, where visual inputs terminate in an “accessary calyx” attached to one side of the fused calyx (Vogt et al., 2016). A similar arrangement involving visual inputs has been described from the swallowtail butterfly (Kinoshita et al., 2015).

Of particular note is the extensive organization of GAD-immunoreactive arborizations that extend into every domain of the *H. nudus* calyces. These provide dense systems of minute processes, the dispositions of which would allow synaptic contact with every neuronal process supplying a microglomerus. Accepting that varunid intrinsic neurons correspond to insect Kenyon cells, these GAD-immunoreactive systems suggest correspondence with wide-field anaxonal inhibitory neurons that extend throughout the *Drosophila* calyx (‘APL neurons’: Liu and Davis, 2009; Amin et al., 2020). Consequently, such putatively inhibitory anaxonal neurons in the crab suggest a global role in local synaptic inhibition at microglomeruli. Arrangements of aminergic neurons in the calyces and columns further correspond to mushroom body input and output neurons recognized in *Drosophila* (Aso et al., 2014; Owald et al., 2015). It is also notable that highly localized and densely branching TH- and 5HT-immunoreactive neurons occur at different levels of the crab’s calyces. These correspond to similar dispositions of immunoreactive neurons in the layered calyces of columnar mushroom bodies, including those of stomatopods and shrimps (Wolff et al., 2017; Sayre and Strausfeld, 2019). Neurons with the same immunoreactive signatures are associated with columns, from their origin at the calyces out to the level of their “fuzzy” interface with gyri and also in the gyri. Their successive levels in a column approximate the arrangements of output neurons from successive partitions of insect mushroom bodies, as exemplified in cockroaches and *Drosophila* (Li and Strausfeld, 1998; Tanaka et al., 2008; Aso et al., 2014b); and in crustaceans by the successive fields of efferent neurons in the mushroom bodies of Stomatopoda, Stenopodidea and Caridea (Wolff et al., 2017; Sayre and Strausfeld, 2018; Strausfeld et al., 2020).

### Is the organization of gyri an outlier, and what is its homologue in other lineages?

Comparisons of the varunid brain with those of other pancrustaceans meet an impasse when considering the lateral protocerebrum’s system of gyri, a cortex-like feature that appears to be special to Brachyura. As described above, there is no obvious terminus of a mushroom body column: its efferent neurons merge with systems of local interneurons in the gyri and, with these local interneurons, they extend amongst the dendrites of efferent neurons that leave the gyri. If these arrangements do not correspond to organization in other pancrustaceans and mandibulates, then what developmental events might have given rise to such a departure from the reptantian ground pattern?

Three developmental studies on the malacostracan lateral protocerebrum should be noted. Two demonstrate early stages of the formation of the larval lateral protocerebrum, in which there is not yet evidence for morphogenetic reorganization in what will become the lateral protocerebrum (Harzsch and Dawirs1996; Sintoni et al., 2007). The third demonstrates that massive reorganization and new generation of neuropils can accompany late development in the pelagic larva (Chan and Cronin, 2018). All of those authors have indicated that crustaceans can undergo radical metamorphosis during the change from a pelagic larva to a preadult, the larval stage providing pools of neural precursor cells for subsequent development. Those precursors would likely contribute not only to the formation and differentiation of adult centers in the lateral protocerebrum but also to any taxon-specific morphogenic rearrangements.

Might the relationship of the insect mushroom body with the rest of the brain suggest an origin for the gyri? In insects, many of the mushroom body efferent neurons terminate in neuropils of the superior medial protocerebrum (honeybee: Strausfeld, 2002; cockroach: Li and Strausfeld 1997, 1999; *Drosophila*: Ito et al., 1998; Tanaka et al., 2008; Aso et al., 2014a,b). This is the most rostral part of the forebrain in insects, comprising dense networks of local interneurons that are interposed between the terminals of mushroom body outputs and the dendrites of interneurons whose axons terminate in successive strata of the central complex’s fan-shaped body (Phillips-Portillo and Strausfeld, 2012). A comparable arrangement in crustaceans with eyestalks requires that efferent neurons (particularly TH-immunoreactive MBONs) from the mushroom bodies extend axons from that distant origin into the midbrain’s superior protocerebrum. These connections are suggested by silver-stained brains of *H. grapsus* showing numerous axons from the eyestalk nerve directly target the medial protocerebrum. This speaks against gyri being medial neuropils displaced distally.

One possible interpretation of the gyri is that they represent enormously expanded versions of tubercles, swollen parts of a column typifying mushroom bodies of certain caridid shrimps (in Stenopodidae and Thoridae: Sayre and Strausfeld 2019) and insects (Zygentoma, Ephemeroptera: Farris, 2005). Tubercles of caridid (or insect) mushroom bodies are compact, comprising tangled extensions of intrinsic neuron processes associated with a concentrated system of TH- and 5HT-immunorecative processes intersecting that level of the column. Those arrangements are absent from gyri. Although there are TH- and 5HT-immunorecative fields in gyri, these do not appear to interact with intrinsic fibers in the columns. Nevertheless, that gyri are as strongly DC0-immunoreactive as the calyces and columns suggests that they play crucial roles in memory acquisition.

Far more elaborated than a tubercle, gyri comprise many morphologically distinct types of neurons, including efferents of unknown antigenicity. The presence in gyri of local interneurons and rectilinear networks further indicates an identity that is indirectly associated with the mushroom body. A clue is suggested by other Reptantia, in which certain mushroom body outputs target neuropils at deep levels of the lateral protocerebrum (Mellon et al., 1992a,b; McKinzie et al., 2003). If these less well-studied neuropils became inverted, they would overlie the inverted brachyuran mushroom bodies as do the gyri (Figure 12d,e). In that scenario, the relationship between a mushroom body and its target neuropils is essentially the same as in other reptantians, except a consequence of inversion is that those neuropils have been able to undergo unparalleled lateral expansion and gyrification.

### The varunid mushroom body in the context of pancrustacean evolution and cognition

The organization of characters defining mushroom bodies is phenotypically identical in stomatopods and insects (Wolff et al., 2017). The same characters have since resolved divergent homologues in all malacostracan crustaceans except Brachyura, the most recent lineage of decapod crustaceans (Wolfe et al., 2019). By identifying and describing the mushroom body of the shore crab, the present study completes the recognition of mushroom body diversification and phenotypic correspondences, thereby enabling future transcriptomic verification or rejection.

In considering the possible significance of the varunid mushroom body being larger and more elaborate than that of any other pancrustacean it is relevant to consider a well-studied proxy that shares with mushroom bodies circuit and genetic correspondences, as well as cognitive properties. For example, it has been suggested that the cerebellum, caudal to the vertebrate midbrain-hindbrain boundary, would be a fitting proxy because of comparable arrangements of parallel fibers (Schürmann, 1987; Farris, 2011, Li et al., 2020). However, mushroom bodies are situated rostral to the arthropod brain’s deutocerebral-tritocerebral boundary. A homologous location is occupied by the hippocampi, rostral to the vertebrate midbrain-hindbrain boundary (Bridi et al., 2020). Mushroom bodies and hippocampi are restricted, respectively, to the protocerebrum and its vertebrate homologue the telencephalon (Hirth and Reichert, 1999). Neuroanatomical arrangements that are intensely immunoreactive to antibodies against DC0 are shared by the hippocampus and mushroom bodies, as are at least sixteen orthologous genes required for the same functions in both (Wolff and Strausfeld, 2016). Gene expression profiling of early mushroom body development in *Drosophila* and of the murine pallium resolve further correspondences (Tomer et al., 2010).

A range of behaviors supported by both the insect mushroom bodies and hippocampus relate to allocentric memory, recall of place, and the use of space (Krebs et al., 1989; López et al, 2003; Salas et al., 2003; LaDage et al., 2009; Montgomery et al., 2016; van Djik et al., 2017). Insects that are permitted explorative foraging acquire enlarged mushroom bodies compared with constrained siblings, a property also pertaining to the hippocampus (Montgomery et al., 2016; van Djik et al., 2017; Basil et al, 1996). As is true for mammalian and avian hippocampi voluminous mushroom bodies indicate a species’ reliance on spatial or social cues (Healy and Krebs, 1992; Withers et al., 1993, 2008; Ott and Rogers, 2010; Molina and O’Donnell, 2007, 2008). Hippocampal lesions impair place memory (Day et al., 2001; Clark et al., 2005), as do lesions of mushroom bodies (Mizunami et al., 1998; Buehlmann et al., 2020; Kamhi et al., 2020). The volume and neuronal complexity of mushroom bodies in insects also relates to species with highly developed spatial cognition (cockroaches, hymenopterans; Strausfeld et al., 2009). The enormous mushroom bodies of varunid crabs suggest comparable cognitive properties relating to space. Shore crabs are opportunistic generalists that live at the interface of two biotopes, marine and terrestrial. They are adept at learning complex tasks (Tomsic and Romano, 2013), such as maze learning (Davies et al., 2019), organizing “ad hoc” collaborative actions, establishing social status (Tanner and Jackson, 2012; Kaczer et al., 2007), and memorizing acquired motor skills (Hughes and O’Brien, 2001). Operant conditioning and the retention of contextual memories suggest considerable intelligence (Abramson and Feinman, 1990; Pereyra et al., 1999).

The present description of the varunid mushroom body concludes studies establishing that divergent evolution of the crustacean mushroom body maps to specific malacostracan lineages. These findings offer hitherto unexplored opportunities for relating divergent cognitive centers to specific ecologies and behavioral repertoires required to negotiate them.

## Materials and Methods

### Animals

Crabs (*Hemigrapsus nudus*), with carapace widths between 6 and 8 cms, were collected from designated sites on San Juan Island, Washington, affiliated with the Friday Harbor Marine Biology Laboratory (University of Washington, Seattle). Two further shipments of living crabs were sent to the Strausfeld laboratory and maintained at 15°C in tanks containing seaweed continuously moistened with salt water. A total of 76 animals were used for this study over an 8-year period. Twenty were used for immunohistology, 8 for osmium-ethyl gallate staining, 10 for Bodian reduce silver staining and 38 for Golgi impregnations.

### Neuropil staining with osmium-ethyl gallate

A modified ethyl gallate method (Wigglesworth 1957) was used to stain entire brains that were then embedded in plastic and serial sectioned. These were used for Amira 3D reconstructions. After cooling to complete immobility, the eyestalks were opened to expose the lateral protocerebrum. The head was then immediately sliced off behind the brain and with the eyestalks immersed in 0.13 M cacodylate buffered (pH 6.8; Fisher Scientific, Cat#: 50-980-232) 1% paraformaldehyde with 2% glutaraldehyde (Electron Microscopy Sciences, Hatfield, PA). The rest of the animal was placed in a −40°C freezer. After overnight fixation undissected tissue was soaked in buffer. The lateral protocebra and midbrain were then freed of overlying exoskeleton. Next, tissue was placed for 2 h in 0.5% osmium tetroxide in cacodylate buffer, continuously rocked and kept at around 4°C in the dark. After a washing in cold buffer, tissue was further well washed in distilled water while raising the temperature to 20°C. Tissue was then immersed in aqueous 0.5% ethyl gallate with constant rocking. Tissue was periodically checked and when dark blue (after about 30 minutes) thoroughly washed in distilled water, dehydrated, and embedded in Durcupan to be cut serially into 15μm thick serial sections using a sliding microtome.

### Neuropil staining with reduced silver

Bodian’s original method (Bodian, 1936) was used, after fixing tissue in an admixture consisting of 75 parts absolute ethanol, 5 parts glacial acetic acid, 20 parts 16% paraformaldehyde (Electron Microscopy Sciences; Cat# 16220; Hatfield, PA). As for the osmium-ethyl gallate method, crabs were cooled to complete immobility, the relevant parts removed, opened and fixed in AAF (70 parts absolute ethanol, 5 parts glacial acetic acid, 25 parts 10% E. M. grade formaldehyde). After 2-4 hours neural tissue was freed under 70% ethanol, then dehydrated to absolute ethanol before clearing in terpineol. It was next soaked in Xylol raising the temperature to 65°C. Finally, tissue was transferred, via a mixture of 50:50 Xylol-Paraplast, into pure Paraplast (Sherwood Medical, St. Louis, MO) held at 65°C in shallow aluminum cookie dishes. After three changes, each about 3-4 minutes long, the final dish was placed on icy water for the Paraplast to immediately begin to solidify. After serial sectioning at 10μm, sections were mounted on glycerin-albumin-subbed slides, dried, and dewaxed. Hydrated material was incubated at 60°C in 2-5% silver proteinate (Roques, Paris) for 24 h with the addition of 1-4 g pure copper shot/100 ml. Afterward, sections were conventionally treated with hydroquinone, gold toned, fixed, and mounted under Permount (Fisher, Springfield, NJ).

### Silver chromate (Golgi) impregnation

Animals were anesthetized over ice. The midbrain and eyestalk neural tissue was dissected free from the exoskeleton and its enveloping sheath in ice-cold chromating solution comprising 1 part 25% glutaraldehyde (Electron Microscopy Sciences; Cat# 16220; Hatfield, PA), 5 parts 2.5% potassium dichromate (Sigma Aldrich; Cat# 207802; St. Louis, MO) with 3–12% sucrose, all dissolved in high pressure liquid chromatography (HPLC)-grade distilled water (Sigma Aldrich; Ca#270733). Central brains and their detached lateral protocerebra (tissues) were placed in fresh chromating solution overnight, in the dark at room temperature (RT). Next, tissues were briefly rinsed (30 seconds with swirling) in 2.5% potassium dichromate and transferred to an admixture of 2.5% potassium dichromate and 0.01% osmium tetroxide (Electron Microscopy Sciences; Cat# 19150; Hatfield, PA) for 24 hrs. Tissues were again washed in 2.5% potassium dichromate and immersed for 24 hr. in a fresh chromating solution. Before silver impregnation, tissues were decanted into a polystyrene weighing dish, swirled in 2.5% potassium dichromate and then gently pushed with a wooden toothpick into a glass container containing fresh 0.75% silver nitrate (Electron Microscopy Sciences; Cat# 21050; Hatfield, PA) in HPLC water. Tissues were twice transferred to fresh silver nitrate, then left overnight in silver nitrate. Throughout, metal was not allowed to come into contact with tissue. Tissues were rinsed twice in HPLC water. To achieve massive impregnation of neurons, after treatment with silver nitrate tissues were washed in HPLC water and again immersed in the osmium-dichromate solution for 12hrs. and then transferred to silver nitrate as described above. After washing in HPLC water and dehydration through a four step ethanol series from 50% to 100% ethanol tissues were immersed in propylene oxide (Electron Microscopy Sciences; Cat# 20401; Hatfield, PA), and left for 15min. They were then subject to increasing concentrations of Durcupan embedding medium (Sigma Aldrich; Cat# 44610; St. Louis, MO) in propylene oxide during 3-5 hours. Tissues were left overnight at RT in pure liquid Durcupan and then individually oriented in the caps of BEEM capsules. These were then married to the inverted capsule cut open at its base, and were topped–up with Durcupan. Filled capsules were placed in a 60°C oven for 18–24 hr. for plastic polymerization. Cooled blocks were sectioned at a thickness of 40-50μm (max) and mounted using Permount mounting medium (Fisher Scientific; Cat# SP15-100; Hampton, NH).

### Immunohistochemistry, antibodies

Visualization of arrangements of immunostained neurons that further define mushroom bodies was achieved using antibodies listed in Table 1. An antibody against DC0 that recognizes the catalytic subunit of protein kinase A in D. melanogaster (Skoulakis et al., 1993) is preferentially expressed in the columnar lobes of mushroom bodies across invertebrate phyla, including members of Lophotrochozoa (Wolff & Strausfeld, 2015). Western blot assays of DC0 antibodies used on cockroach, crab, centipede, scorpion, locust, remipede, and millipede neural tissue reveal a band around 40 kDa, indicating cross-phyletic specificity of this antibody (Stemme et al., 2016; Wolff & Strausfeld, 2015). Antibodies against synapsin (Klagges et al., 1996) and α-tubulin (Thazhath et al., 2002) were used, often in conjunction with other primary antibodies, to identify dense synaptic regions and general cellular connectivity. Both antibodies likely recognize highly conserved epitope sites across Arthropoda and have been used previously in crustaceans (Andrew et al., 2012; Brauchle et al., 2009; Harzsch et al., 1997; Harzsch & Hansson, 2008; Sullivan et al., 2007). Antibodies against serotonin (5HT), glutamic acid decarboxylase (GAD), and tyrosine hydroxylase (TH) were used in this study to describe neuronal organization and in distinguishing neuropil boundaries. 5HT is an antibody that has proven to be invaluable for neuroanatomical studies across Arthropoda (Antonsen & Paul, 2001; Harzsch & Hansson, 2008; Nässel, 1988). Previous studies have used 5HT as a comparative tool for neurophylogenetic analysis (Harzsch & Waloszeck, 2000). Antibodies against GAD and TH were used to detect the enzymatic precursors of gamma aminobutyric acid (GABA) and dopamine, respectively. These two antibodies do not require the use of alternative fixation methods, making them compatible with synapsin and α-tubulin labeling, while avoiding the need to use glutaraldehyde. Comparisons of anti-GAD and anti-TH immunolabeling with that of their respective derivatives have demonstrated that these antibodies, respectively, label putative GABAergic and dopaminergic neurons (Cournil et al, 1994; Crisp et al., 2001; Stemme et al., 2016; Stern, 2009).

### Immunohistochemistry, application

Methods follow those used for two recent studies on the eumalacostracan brain (Sayre & Strausfeld, 2019; Strausfeld et al., 2020). Animals were anesthetized to immobility using ice. Brains detached from nerve bundles leading to the eyestalks were first dissected free and the eyestalks were then removed. Immediately after removal, tissue was immersed in ice-cold fixative (4% paraformaldehyde in 0.1 M phosphate-buffered saline (PBS) with 3% sucrose [pH 7.4]). Midbrains and lateral protocerebra with their intact optic lobes were desheathed and left to fix overnight at 4°C. Next, tissue was rinsed twice in PBS before being transferred to the embedding medium (5% agarose with 7% gelatin) for 1 hr. at 60°C before cooling to room temperature in plastic molds. After solidification, blocks were removed from the molds and postfixed in 4% paraformaldehyde in PBS for 1 hr. at 4°C. The blocks were then rinsed twice in PBS and sectioned at 60 μm using a vibratome (Leica VT1000 S; Leica Biosystems, Nussloch, Germany). Next, tissue sections were washed twice over a 20 min. period in PBS containing 0.5% Triton-X (PBST). Tissue was subsequently blocked in PBST with 0.5% normal donkey serum (NDS; Jackson ImmunoResearch; RRID:AB_2337258) for 1hr before primary antibody incubation. Primary antibodies were added to the tissue sections at dilutions listed in Table 1. Sections were left overnight on a rotator at room temperature. Sections were next rinsed in PBST six times over the course of an hour. Donkey anti-mouse Cy3 and donkey anti-rabbit Cy5 or Alexa 647 (Jackson ImmunoResearch; RRID: AB_2340813; RRID: AB_2340607; RRID: AB_2492288, respectively) IgG secondary antibodies were added to Eppendorf tubes at a concentration of 1:400 and spun for 12 min at 11,000 g. The top 900 μL of the secondary antibody solution was added to the tissue sections, which were then left to incubate overnight at room temperature on a rotator. For F-actin staining, tissue sections were left to incubate in a solution containing phalloidin conjugated to Alexa 488 (Thermo Fisher Scientific; RRID: AB_2315147) at a concentration of 1:40 in PBST for 2–3 days with constant gentle agitation following secondary antibody incubation. To label cell bodies, tissue sections were then rinsed twice in 0.1 M Tris–HCl buffer (pH 7.4) and soaked for 1 hr. in Tris–HCl buffer containing 1:2000 of the nuclear stain, Syto13 (ThermoFisher Scientific; Cat# S7575) on a rotator. Next, sections were rinsed six times in Tris–HCl over the course of 1 hr. before being mounted on slides in a medium containing 25% Mowiol (Sigma Aldrich; Cat# 81381) and 25% glycerol in PBS. Slides were covered using #1.5 coverslips (Fisher Scientific; Cat# 12-544E). To verify secondary antibody specificity, primary antibodies were omitted resulting in complete abolishment of immunolabeling. TH immunolabeling required a modified staining procedure with a shorter fixation time as well as antibody incubation in whole unsectioned tissue. Standard fixation times or sectioning the tissue prior to immunostaining resulted in poor or absent labeling as has been described previously (Cournil et al., 1994; Lange & Chan, 2008). For TH labeling, neural tissue was dissected and fixed in 4% paraformaldehyde in PBS containing 3% sucrose for 30–45 min. Following fixation, neural tissue was rinsed twice in PBS, and then twice in 0.5% PBST over the course of 40 min. Tissue was then transferred to blocking buffer containing 5% NDS in 1% PBST and left to soak for 3 hr. TH primary antibody was next added to the blocking buffer at a concentration of 1:250. To assist antibody permeation, whole tissues were microwave-treated for 2 cycles of 2 min on low power (~80 W) followed by 2 min no power under a constant vacuum. Tissue was subsequently left to incubate in primary antibody solution for 2–3 days and was microwave-treated each day. After primary antibody incubation, tissue was washed with 0.5% PBST six times over the course of 2 hr. and then transferred to a solution containing 1:250 Cy3 secondary antibody. Whole mounts were left in secondary antibody overnight on a gentle shaker before being sectioned and mounted as described above. In dual labeling experiments, sectioned tissue labeled with anti-TH was then stained with an additional primary and secondary antibody also as described above.

### Imaging

Confocal images were collected as Tiff files using a Zeiss Pascal 5 confocal microscope (Zeiss; Oberkochen, Germany). Image projections were made using Zeiss Zen System software. Light microscopy images were obtained using a Leitz Orthoplan microscope equipped with Plan Apochromat oil-immersion objectives (X40, X60, and X100). Series of step-focused optical sections (0.5–1.0 μm increments) were collapsed onto a single plane using Helicon Focus (Helicon Soft; Kharkov, Ukraine). Images were transferred to Adobe Photoshop (Adobe Systems, Inc; San Jose, CA) and processed using the Photoshop camera raw filter plug-in to adjust sharpness, color saturation and luminance, texture and clarity.

#### 3D-reconstruction

Serial sections of osmium-ethyl gallate stained eyestalks were imaged using a brightfield Leitz Orthoplan microscope at a pixel scale of 0.5 μm x 0.5 μm. The sections were cut at a thickness of 15 μm using a rotating microtome. To account for compression and error using dry lenses because of the difference in refractive indices between glass/section and air, image sections were digitally adjusted to a voxel size of 0.5 μm x 0.5 μm x 24 μm. Additionally, to obtain a better Z-resolution, two snapshots at two focus planes were taken for each section. The resulting images were manually aligned using non-neuronal fiduciaries in the software TrakEM2 (Cardona et al., 2012).

Neuropils, cell bodies, and large nerve tracts were segmented using the software Amira (Amira 2019.4; Thermo Fisher Scientific; Waltham, MA; USA). We achieved this using the Segmentation Editor, a tool which enables tracing of image data by assigning voxels to materials (i.e. user defined objects such as neuropils, cell bodies, etc.) using a paintbrush tool. Selected sections were outlined in the XY plane and interpolated across sections. Then, using the aid of the interpolated outlines, selected sections were traced in the XY, YZ and XZ planes to create a scaffold of the region of interest. The function “wrap” was then used to interpolate a 3D reconstruction of the object constrained by the scaffold. For visualization purposes, the label field containing “wrapped” labels was interpolated in Z using the “Interpolate Labels” module, adding an additional three slices for each already existing slice while maintaining the overall dimension of the reconstruction. This helped to account for the anisotropic Z-resolution (i.e. “staircase effect”). The 3D model was then visualized using the “Surface Generator” module, which created a 3D polygonal surface mesh. Images used for the figures were generated using the snapshot function. The globuli cell population was estimated on the basis of spherical globuli cells, each with the average radius of 3μm, occupying a volume of 2.5074^3^μm^3^ calculated from the serially reconstructed globuli cell mass.

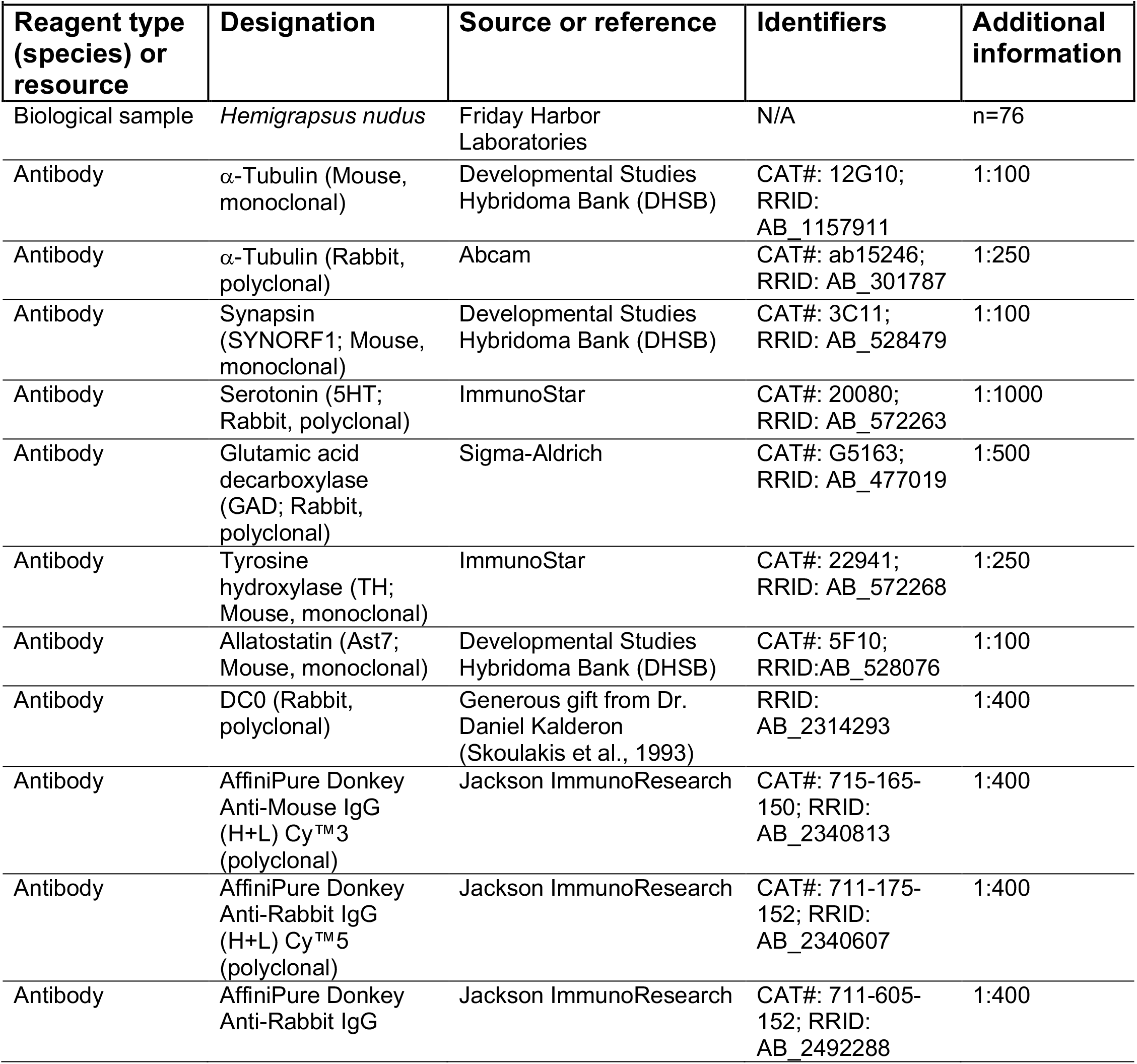
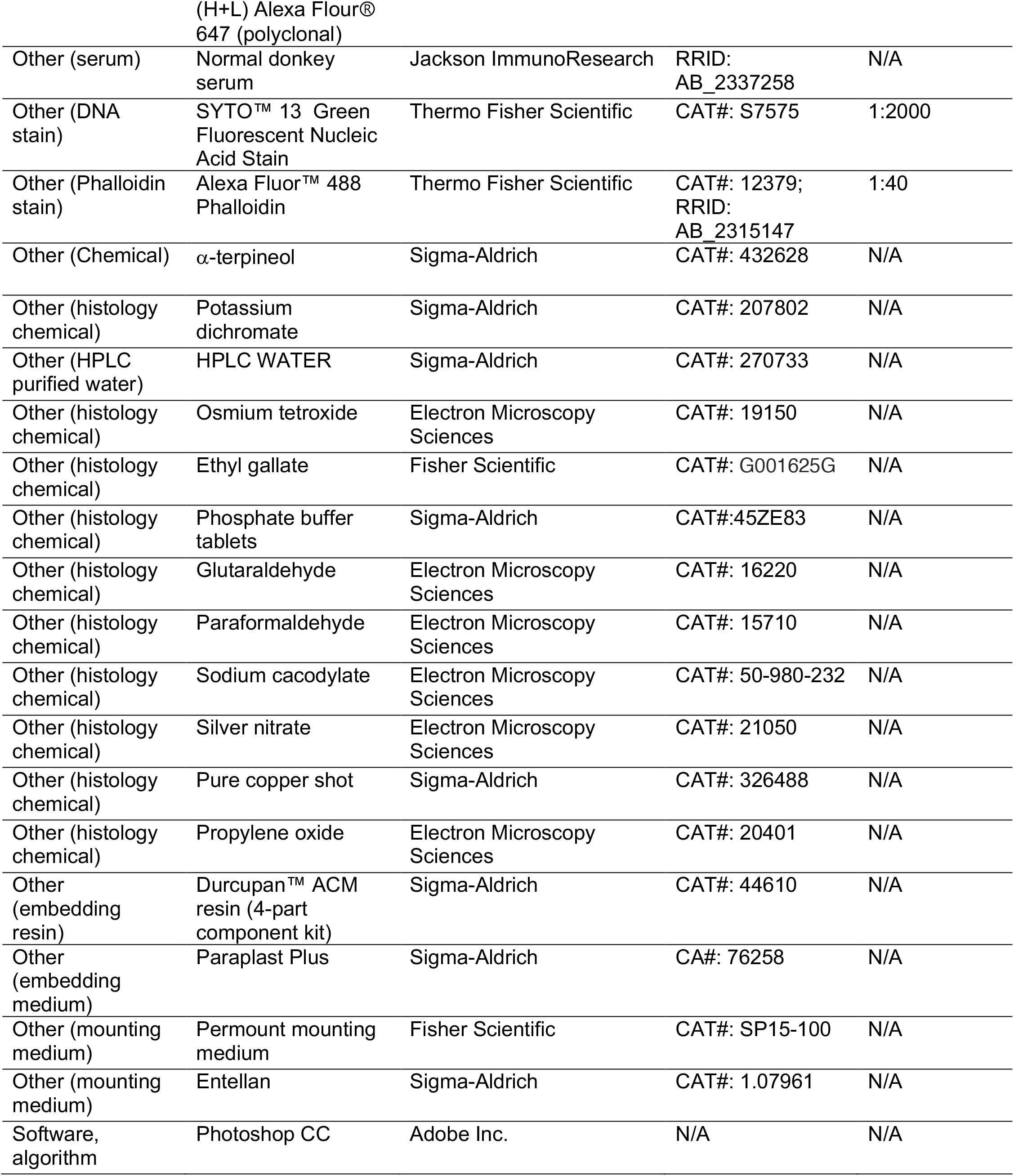
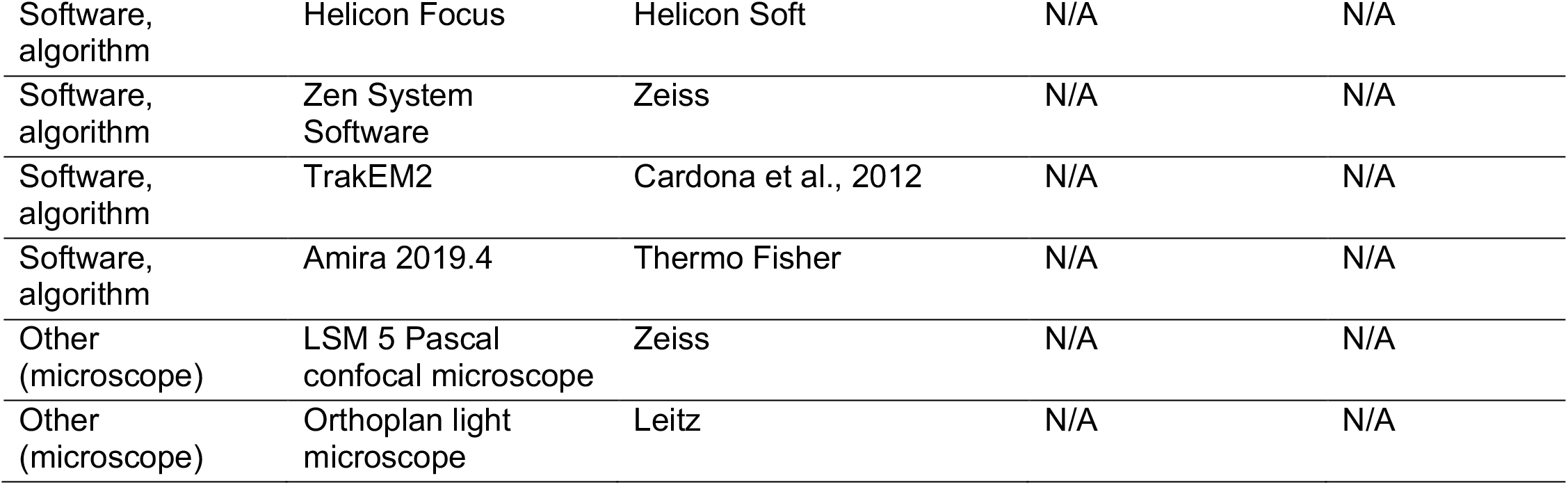
Key Resources Table.

## Terminology

Terms and abbreviations follow recommendations by the insect brain consortium regarding those jointly applicable to insect and crustacean (Pancrustacea) central nervous systems. See Ito et al. (2014). Other terms used for crustacean brain centers follow those of Bellonci (1882) and Sandeman et al. (1993). Other terms are defined in the body of this paper.

## Acknowledgements

The research described here is supported by the National Science Foundation under Grants No. 1754798 awarded to NJS. Our gratitude is once again directed to Daniel Kalderon, Columbia University, New York, for supplying the DC0 antibodies, as he has during the last decade and longer. We thank the staff of the University of Washington’s Friday Harbor Marine Laboratories, San Juan, for their unfailing help in obtaining living specimens. Briana Olea-Rowe and Hannah Joy Miller provided expert technical assistance. We have profited from discussions and advice from Wulfila Gronenberg (University of Arizona) and Frank Hirth (Kings College, University of London) and are indebted to Camilla Strausfeld for her advice, critically discussing versions of the manuscript, suggesting many improvements and meticulously editing the text.

## FIGURE SUPPLEMENTS

**Figure 1 ––– figure supplement 1.**
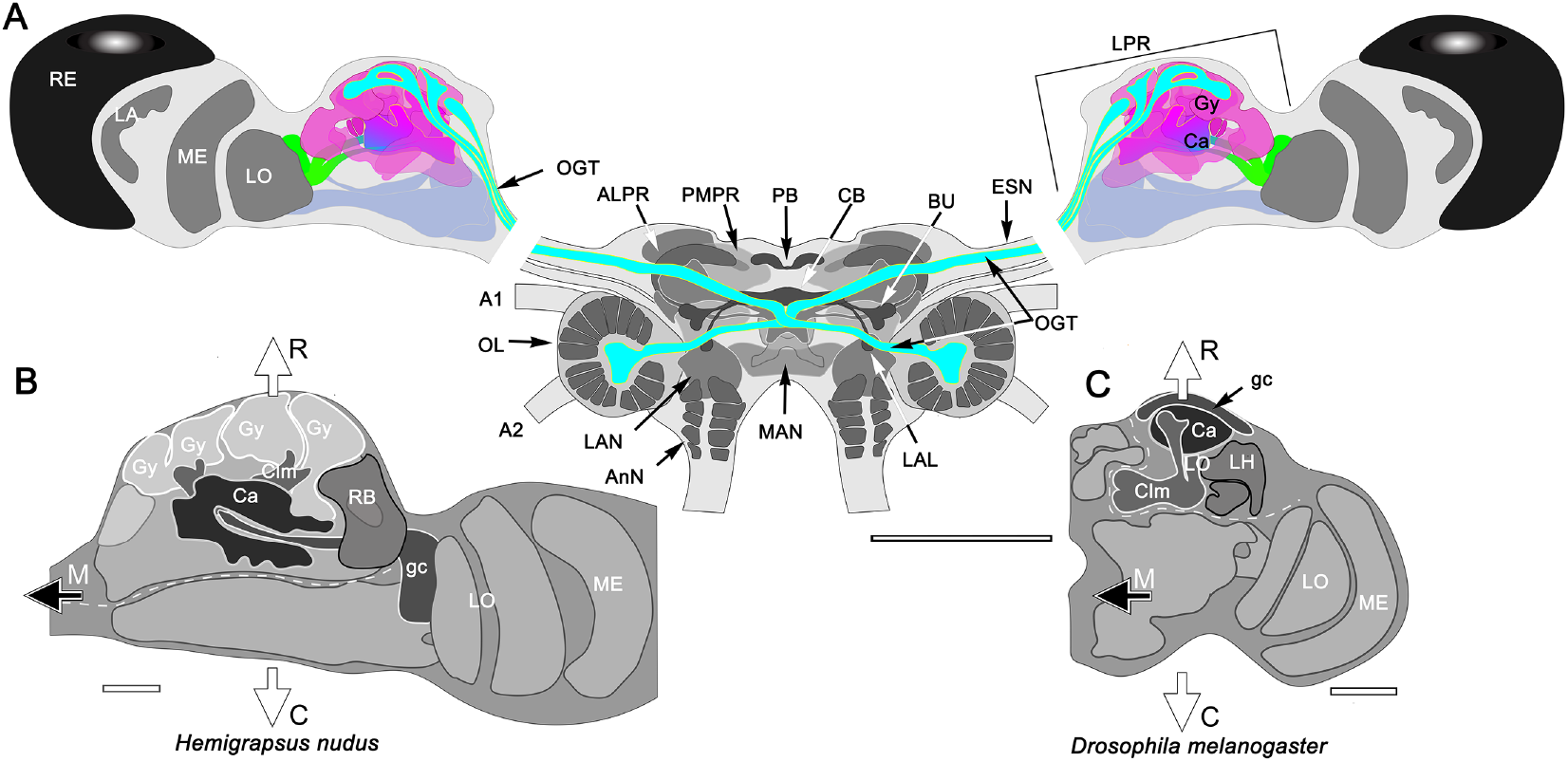
**(a)** Schematic of the *Hemigrapsus* brain (frontal view) and major neuropils. Abbreviations: ALPR, anterior lateral protocerebrum; A1, antennule nerve; A2, antennal nerve; AnN, antenna 2 neuropil; BU, lateral bulb of the central complex; CB, central body; ESN, eyestalk nerve; LAL, lateral accessory lobe; LAN, lateral antennular lobe neuropil; LA, lamina; LO, lobula; LPR, lateral protocerebrum; MAN, medial antenna 1 neuropil; MB, mushroom body; ME, medulla; OL, olfactory lobe; OGT, olfactory globular tract; PMPR, posterior medial protocerebral neuropils; PB, protocerebral bridge; RE, compound retina. The eyestalk nerve ESN) carries *all* axons connecting the brain with the lateral protocerebrum. (**b, c**) Orientations of the varunid lateral protocerebrum aligned with that of *Drosophila*. The white dashed lines in *H. nudu*s divides its rostral lateral protocerebral volume from its caudal volume; in *Drosophila* the dashed line divides the rostral volume of the protocerebrum from the rest of the hemi-brain. In both taxa, the filled arrow M indicates medial, and the open arrows R, C indicate, respectively, the rostral and caudal axes. Both schematics depict a view from the ventral side of the lateral protocerebrum/ hemibrain. Abbreviations: Ca, calyces; Clm, column; gc, globuli cells; Gy, gyri; LH, lateral horn; LO, lobula complex; ME, medulla; RB, reniform body; Scale bars: a, 500μm; b–d 100μm

**Figure 2 ––– figure supplement 1.**
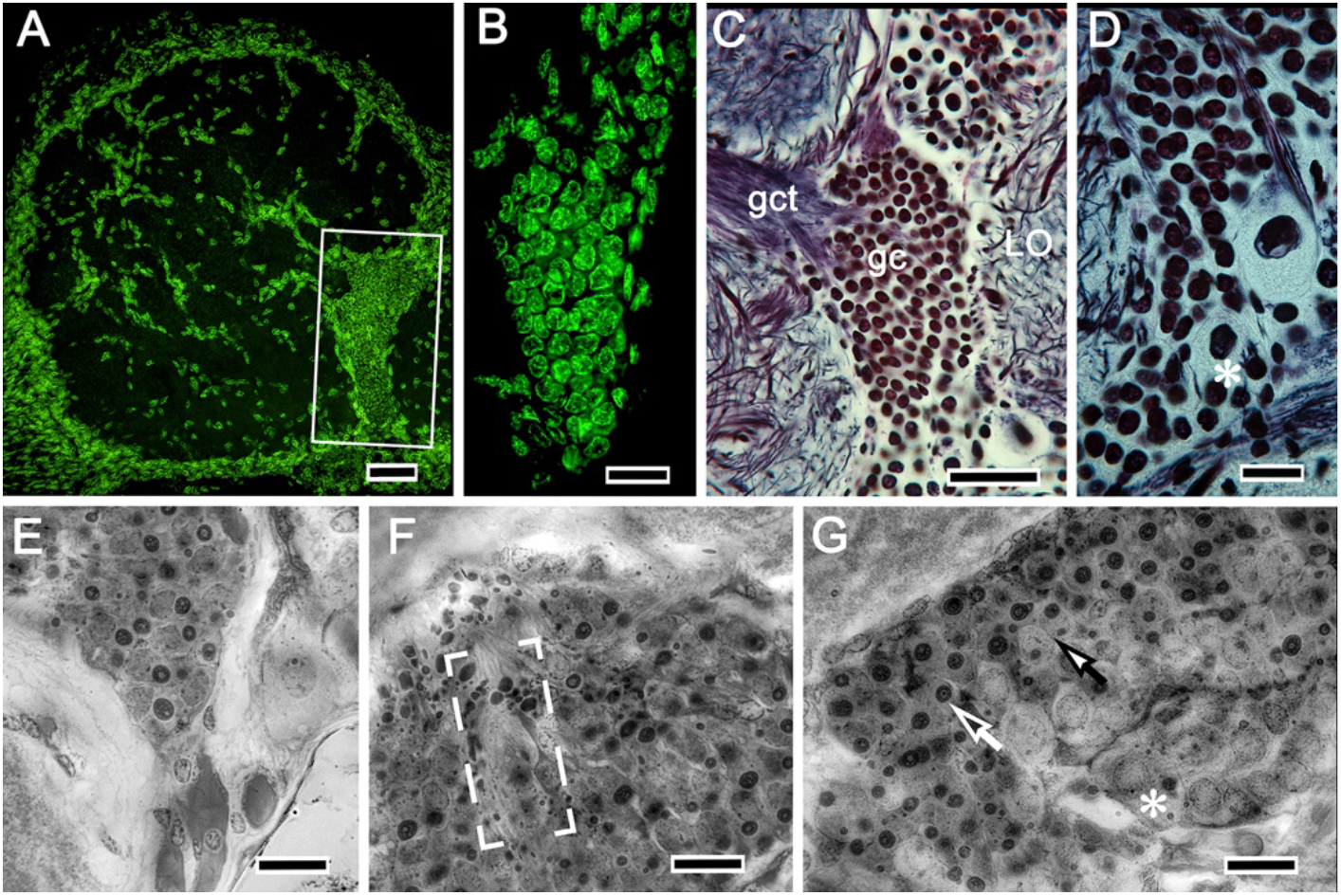
Globuli cells supplying intrinsic neurons to the mushroom bodies. In the varunid crab, globuli cells occupy a unique location between the distal margin of the lateral protocerebrum and proximal margin of the optic lobe as shown in **A**. (**B**) As is typical of globuli cells in other taxa (insects, shrimps) they are minute, here measuring less than 6 μm diameter, the next largest being reniform perikarya as in panel C, upper right. (**D-G**) Like other decapods, crabs molt and grow throughout life, their brains also increasing in size. The globuli cell cluster evidences neuroblasts (asterisk in panel D), and grouped globuli cells suggesting clonal siblings (**E**). Ganglion mother cells are also arranged as aligned groups (**F**) or, as in panel **G**, dividing (closed arrow) between neuroblasts (asterisk) and fully differentiated perikarya (open arrow). Scale bars, A, 50mm; B, 20μm; C, 50μm; D-G, 20μm.

**Figure 2 ––– figure supplement 2.**
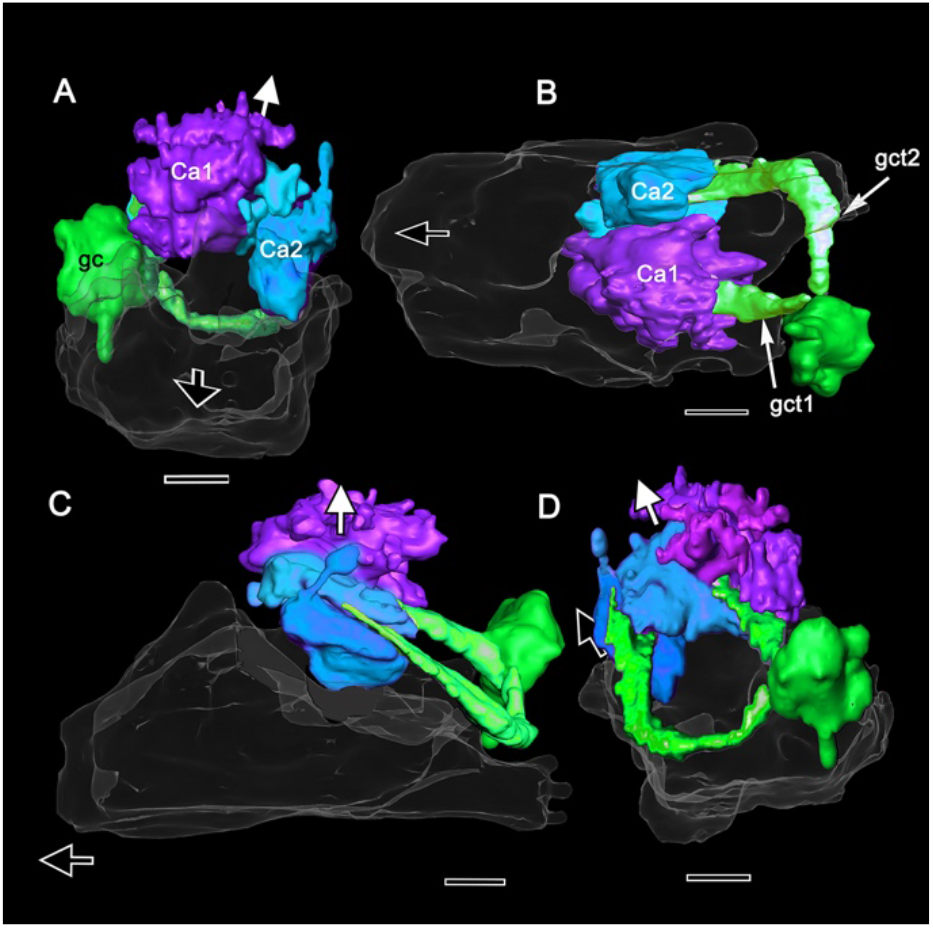
Globuli cells provide two tracts, one to each calyx as shown in this Amira reconstruction of a serially sectioned osmium-ethyl gallate-treated brain. Two tracts composed of neurites from 2,000-3,000 globuli cells from a unified cluster (gc) extend to two calyces (Ca1, Ca2), Ca1 is dorsal to and larger than Ca2. (**A**) Viewed from above, with the medial margin of the lateral protocerebrum closest to the observer, the rostral surfaces of each calyx are highly indented with some finger-like extensions reaching upwards. (**B**) viewed from beneath, with the dorsal calyx 1 closest to the observer, the more caudal levels of the calyces are seen to be layered, this distinction between caudal and rostral volumes also shown viewing into the lateral protocerebrum from its ventral surface (C). (**D**) 180° rotation from panel A suggests greater surface elaboration of Ca1 than Ca2.

**Figure 3 ––– figure supplement 1.**
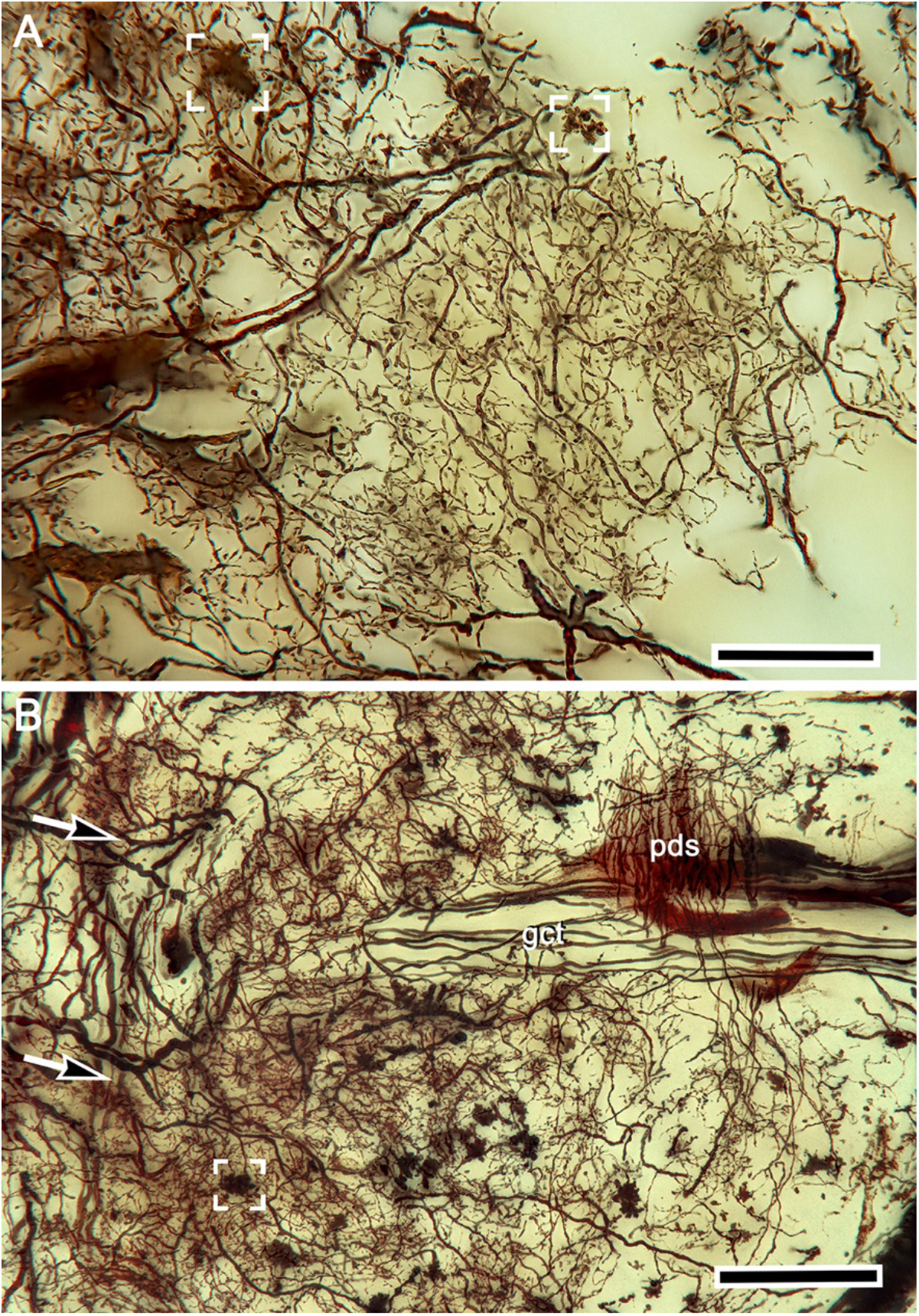
Rectilinear network organization of the calyx and afferent supply. (**A**) Golgi impregnation showing regular network arrangements of intrinsic cell dendrites. Robust afferent terminals are defined by varicose specializations. A single terminal is shown in the box upper right; several convergent terminals in the box upper left. (**B**) The globuli cell tract (gct) extends medially beneath processes belonging to the reniform body’s pedestal (pds). Afferent terminals fibers enter from the left (arrowed) to distribute terminals (one boxed, lower left) amongst densely arranged intrinsic neuron dendrites extending from globuli cell neurites comprising the gct. Scale bars, A, B, 50μm.

**Figure 4 ––– figure supplement 1.**
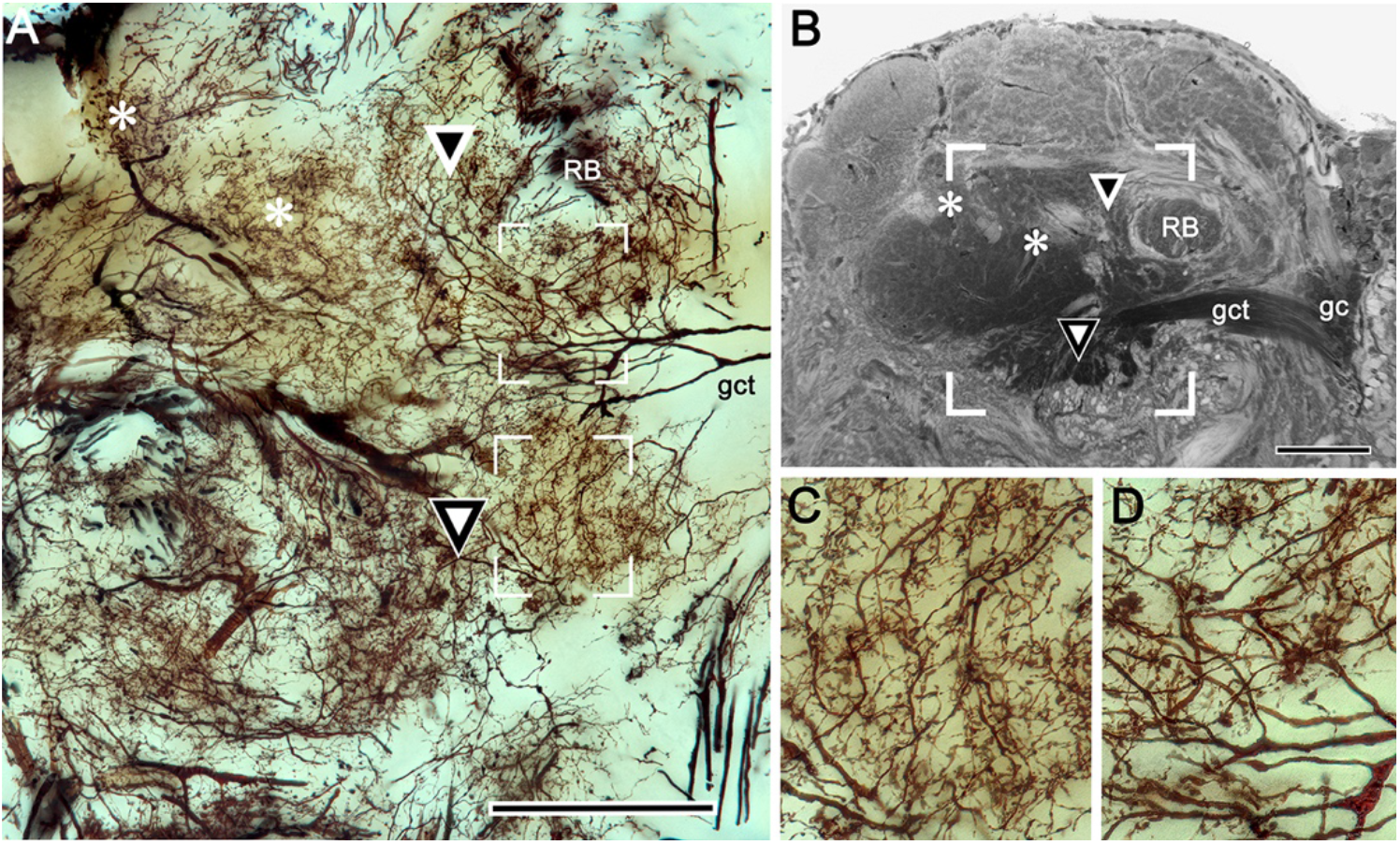
Fidelity of intrinsic neurons fields (**A**) and calycal domains revealed by osmium-ethyl gallate (**B**). The equivalent of the area in panel **A** is indicated by the open box in panel **B**. Corresponding loci in B and A are indicated by symbols. Boxed areas in **A** are enlarged in panels **C**, **D** showing rectilinear organization of intrinsic processes and their convergence at nodes corresponding to microglomeruli. Abbreviations: RB, reniform pedestal; gc, globuli cells; gct, globuli cell tract. Scale bars, A, B, 100μm

**Figure 4 ––– figure supplement 2.**
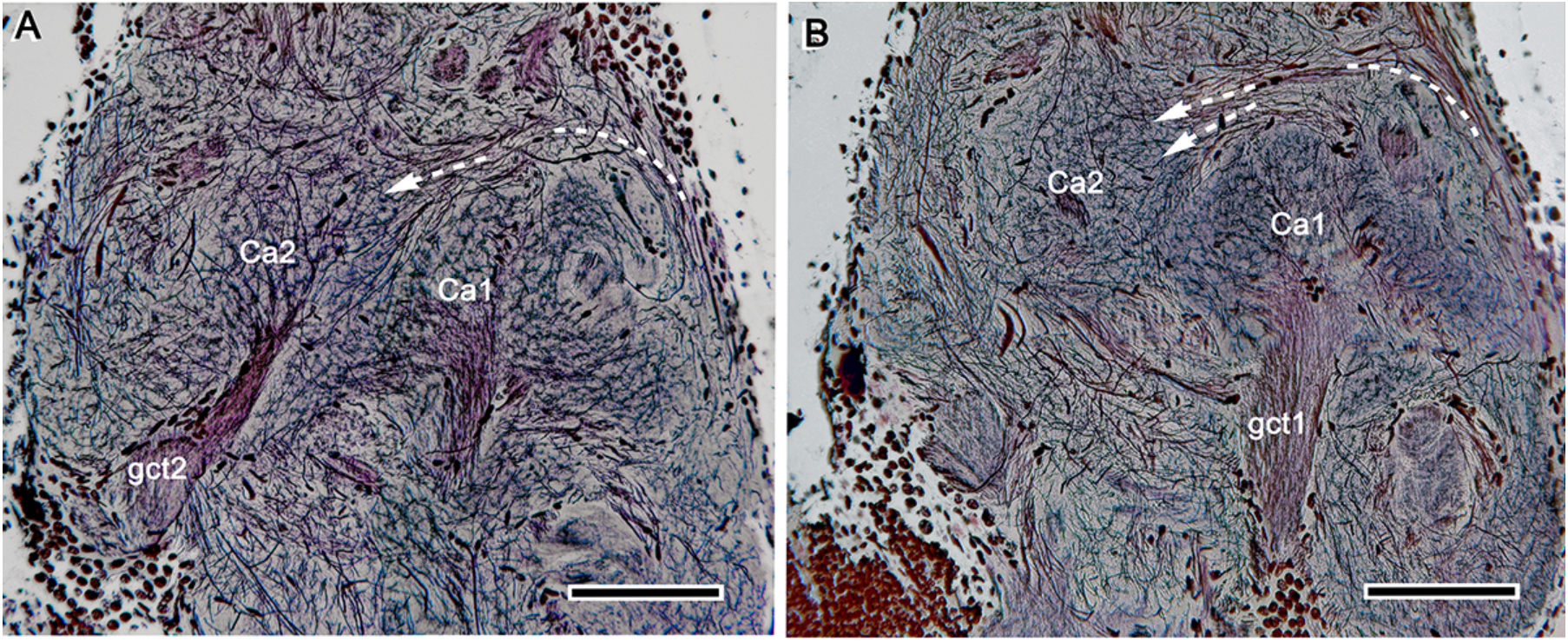
Calyx distinctions. (**A, B**) Two serial silver-stained sections showing the side-by-side disposition of calyces 1, 2 (Ca1, Ca2) each supplied by its own globuli cell tract (gct1, 2). Arrowed paths indicate passage of axons extending to Ca2 from pathways from the optic lobe medulla and reniform body (far right in both panels). Scale bars, 100μm.

**Figure 7 ––– figure supplement 1.**
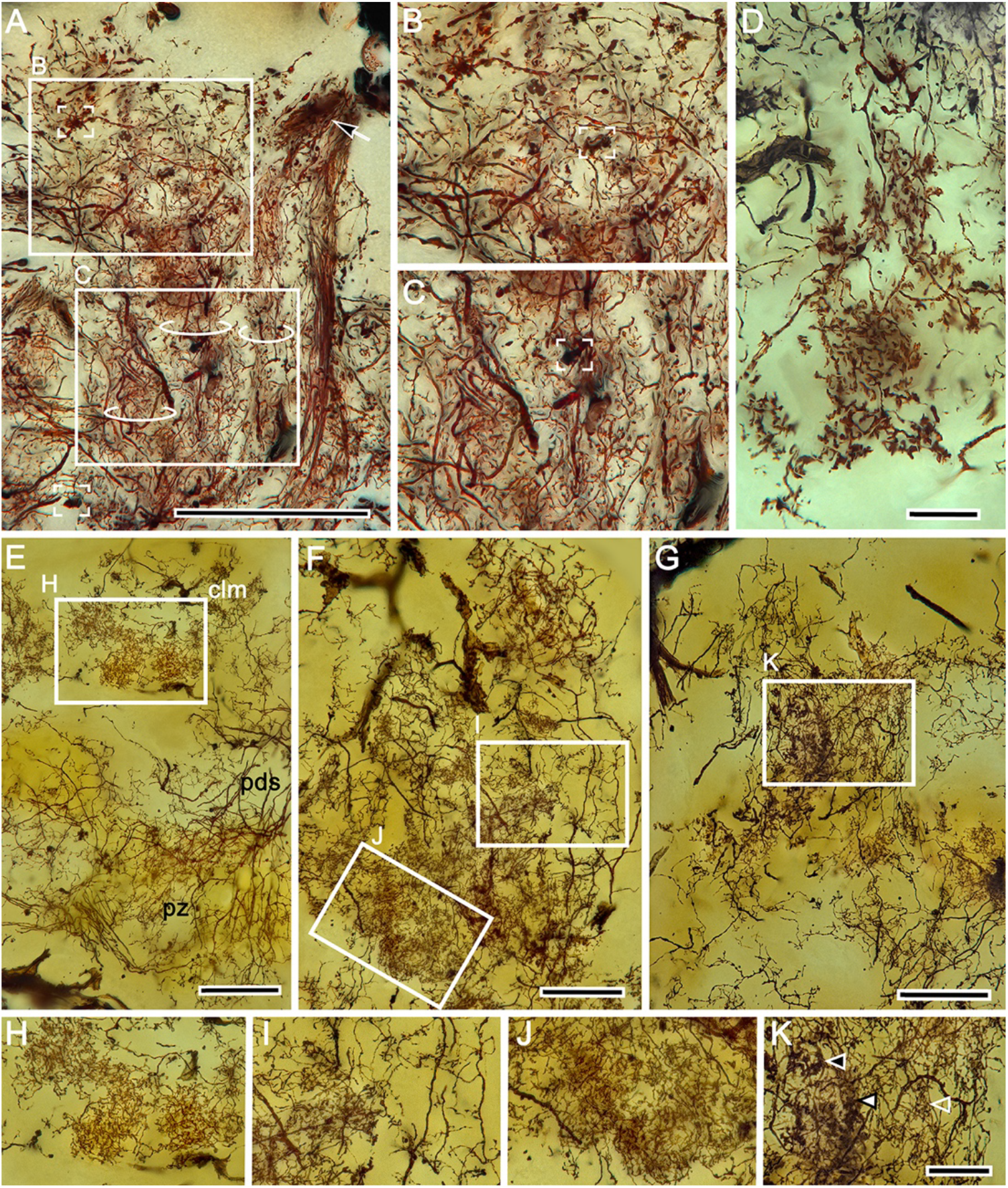
Intrinsic neuron organization in columns. (**A**) Golgi impregnation of a column originating from calyx 2. Three parallel but distinct ensembles of intrinsic neuron processes (circled) all splay outwards immediately beneath the overlying gyriform neuropil, which is here unstained. Details of these ensembles shown in **B**, **C** demonstrate internal complexity. Of note is the presence of bouton-like terminals (two indicated by box brackets) suggesting either afferent supply from the eyestalk nerve or reciprocal inputs from rostral gyri. A cascade arrangement of processes extending parallel to the column from a more rostral level (arrow) may indicate recurrent afferent supply from the gyri. (**D–G**) Columns also contain what appear to be intrinsic elements (**D**) that do not originate from the calyces, as occurs in the mushroom bodies of certain insects (see Discussion). (**E**) There is a clear distinction between the arrangement of processes (enlarged in **H**) in the columns (clm) and the more distributed arrangements in the proximal zone (pz) of the reniform body indicated by its pedestal (pds). (**F**) Further complexity is demonstrated by parallel arrangements of intrinsic neuron and their collaterals suggestive of local circuits (examples indicated in boxed areas in panel F, enlarged in **I**, **J**). (**G**) Dense arrangements near the column’s terminal expansion reveal claw-like specializations each with its particular morphology, within parallel subdivisions of the column (indicated by arrowheads in K). Scale bars, A,100μm; E–G, 25μm; D, H-K, 20μm.

**Figure 7 ––– figure supplement 2.**
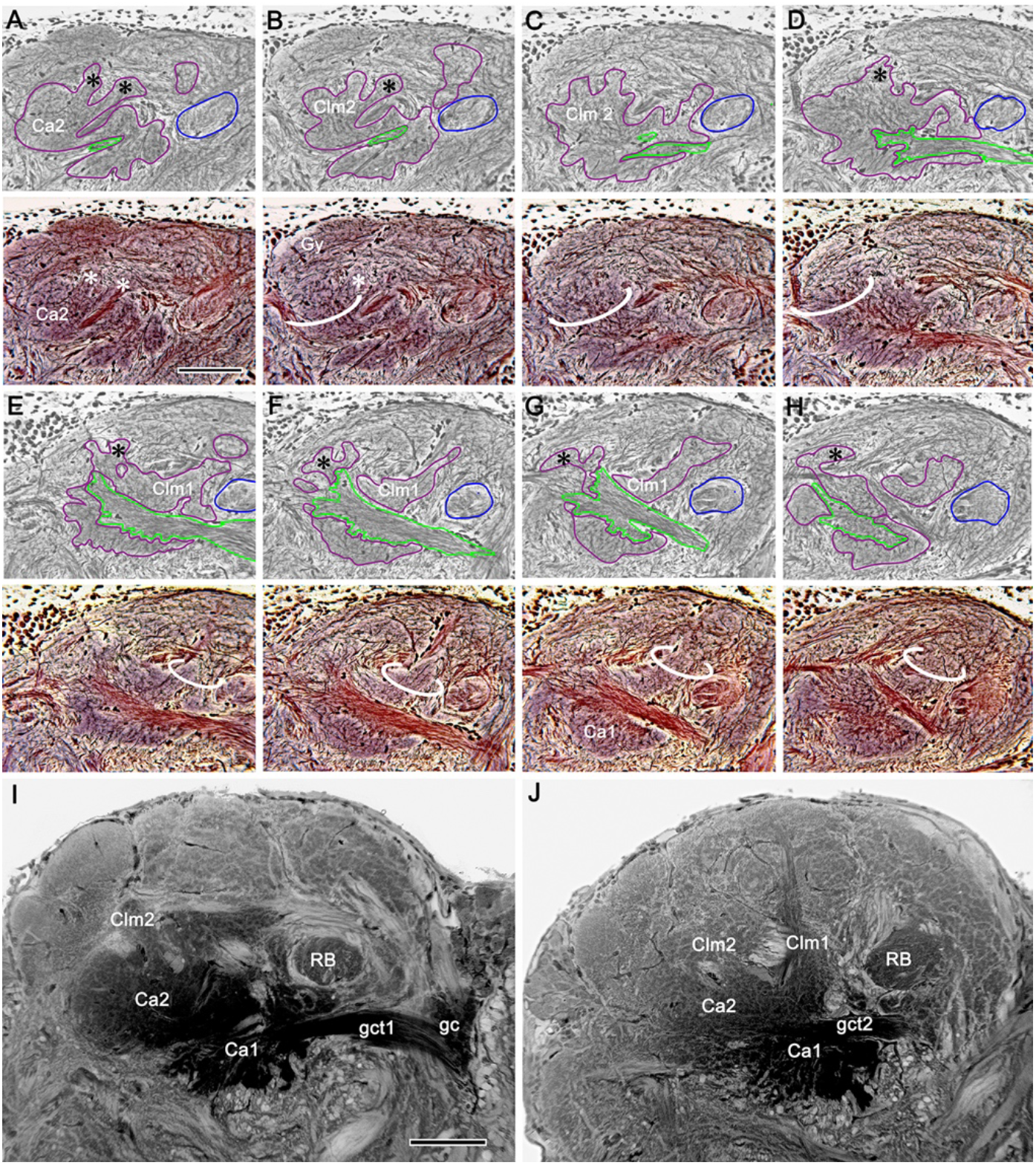
The interface between columns and gyri. (**A-H**) Paired half-tone/color micrographs of selected levels of a silver-stained, serially sectioned RLPR (medial to the left, rostral upwards). Green outlines in panels A-D indicate parts of the globuli cell tract (gct2) to calyx 2 (Ca2); in E-H, parts of the gct1 to calyx 1 (Ca1). Blue outlines indicate cross–sections of the reniform body pedestal (RB); magenta outlines indicate mushroom bodies (calyces and their columns). Identification of the calyces is enabled by their color and microglomeruli as in Ca2, (lower panel A). As in osmium-ethyl gallate sections, the two calyces are too close to identify separately unless viewed downwards from the rostral surface of the LPR (as in Figure 4 - figure supplement 2). Columns (ringed) and smaller finger-like extensions (asterisks) extend outwards to the gyri. Column 2 (Clm2) from calyx 2 (Ca2) is shown in panels A-D. Panels E-H illustrate the origin and width of Clm1 from Ca1. Panels I and J show corresponding arrangements revealed by osmium-ethyl gallate. Scale bars, 100μm.

**Figure 9 –––– figure supplement 1.**
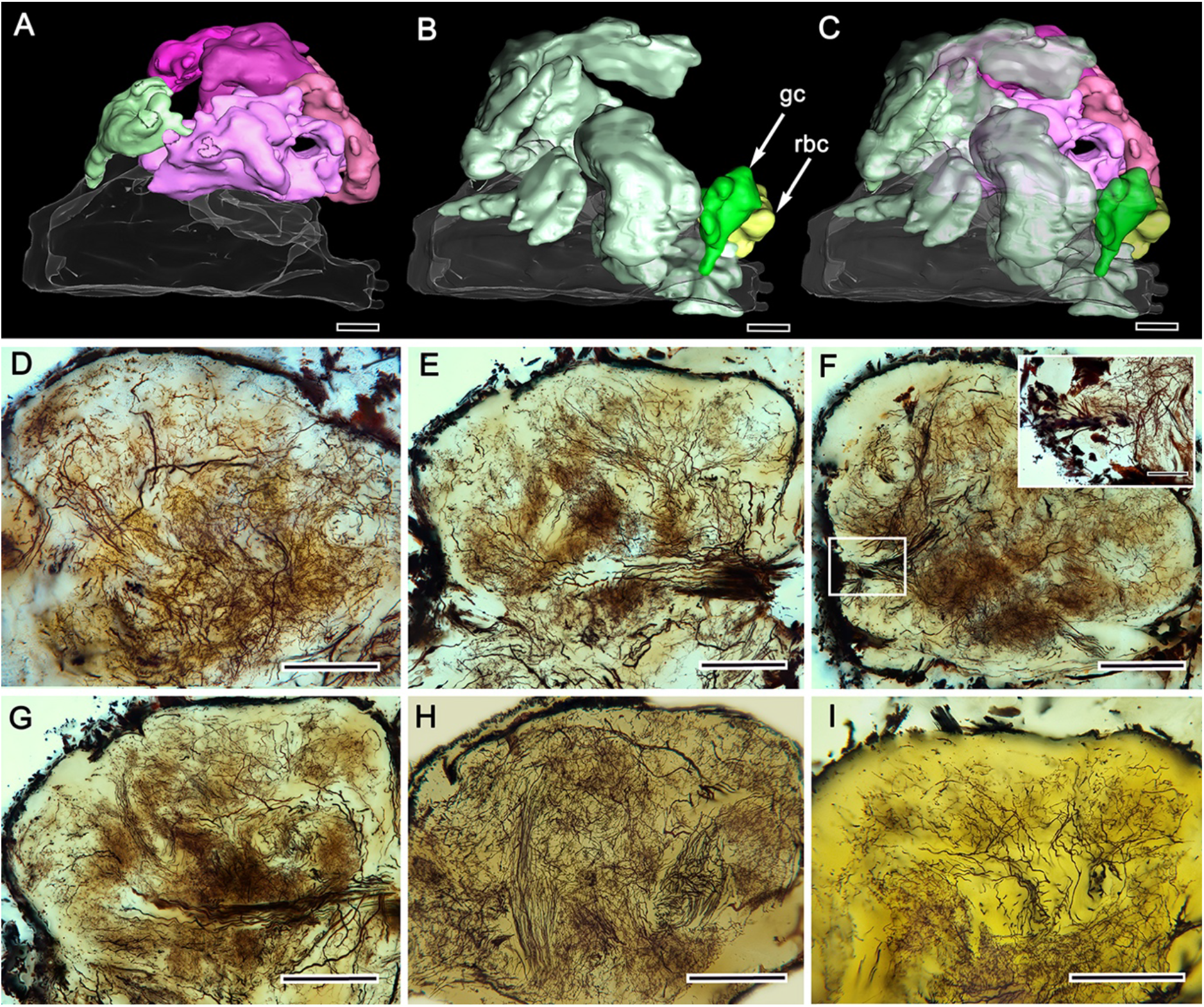
Perikaryal distributions and gyri neurons. (**A–C**). Amira-generated 3-D reconstruction of the gyri in panel A with the proximal gyrus (light green) above the medial root of the eyestalk nerve. Panel B shows extensive patches of neuronal perikarya, with the globuli cell cluster (gc, dark green) and reniform body cluster (rbc, yellow) situated at the distal margin of the RLPR. Merged images in panel C show the relative dispositions of cell body clusters and gyri. (**D-I**) Stitched reconstructions of Golgi mass impregnations resolve top (rostral)-down (F, H), lateral (E, G, I) and sub-gyrus (D) views of neuropils. The boxed area in F includes cell bodies of a lateral patch, enlarged in the upper right inset. Scale bars, A-I, 100μm, inset to panel F, 25μm.

**Figure 10 ––– figure supplement 1.**
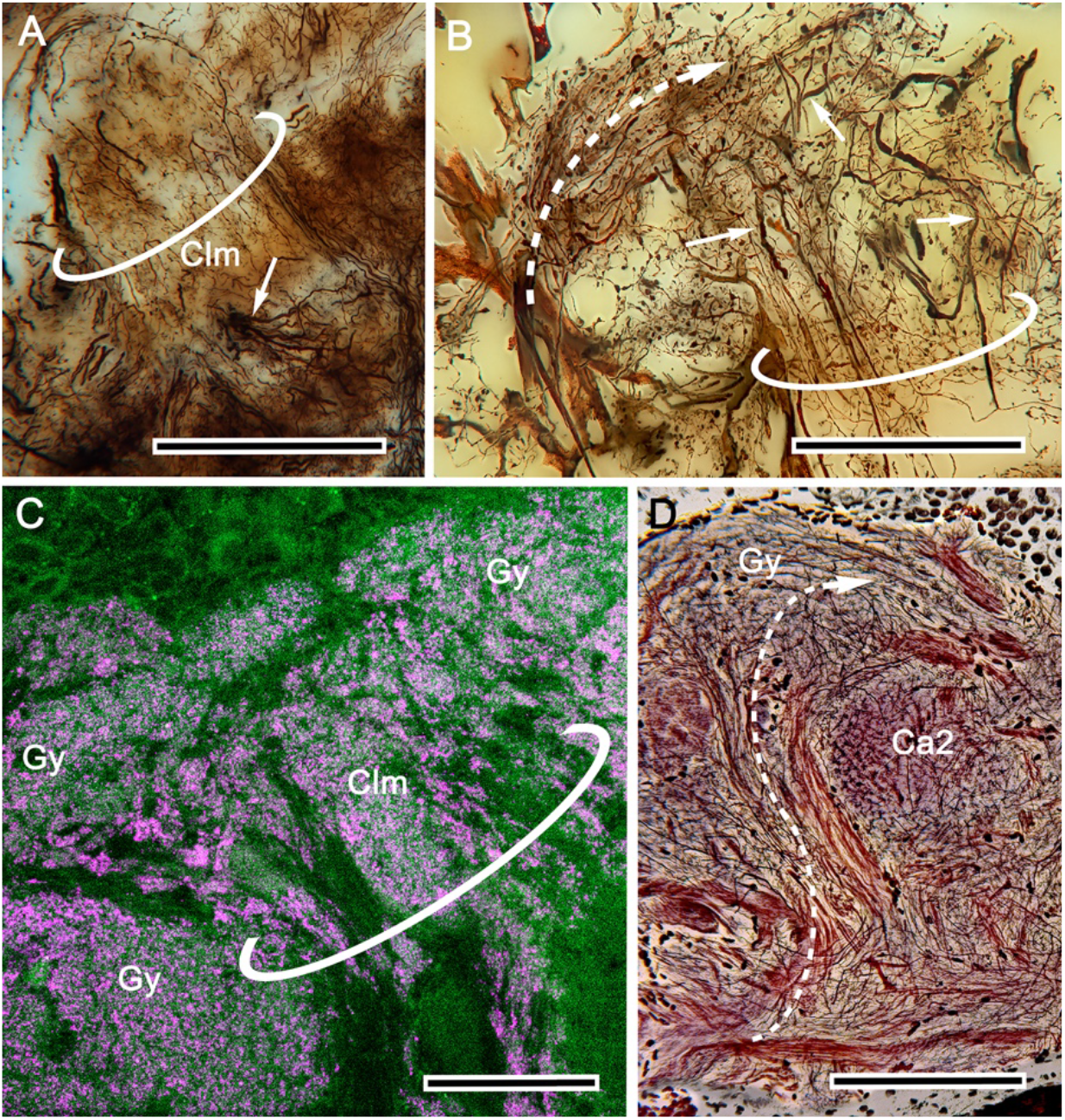
Column-gyrus interface organization.(**A**) Terminus of column 2 from calyx 2 (Clm, circled). The column is notable for its many extremely slender intrinsic processes. Also, visible in this preparation are branches of extrinsic (efferent) neurons extending across the column (arrow). (**B**) Terminus of column 2 (circled) with bundled intrinsic processes and the stouter profiles of efferent neurons (arrowed) ascending from the column into overlying gyral neuropil. This superficial level also receives terminals decorated with small bead-like specializations provided by bundled small-diameter axons from the eyestalk nerve (curved arrow). (**C**) Synapsin (magenta)-actin (green) labelling showing at the terminus of a column (circled) its longitudinal divisions denoted by various sizes and densities of synaptic sites across its width. Similar variation is resolved in flanking neuropil of the gyrus (Gy). (**D**) Reduced silver demonstrates bundles (dashed arrow) from the CLPR ascending outside a column into gyri, demonstrating that multimodal convergence occurs at many levels in the lateral protocerebrum. Part of Ca2 is also visible at this level. Scale bars, A, D, 100μm; B, 50μm; C, 25μm.

**Figure 11 ––– figure supplement 1.**
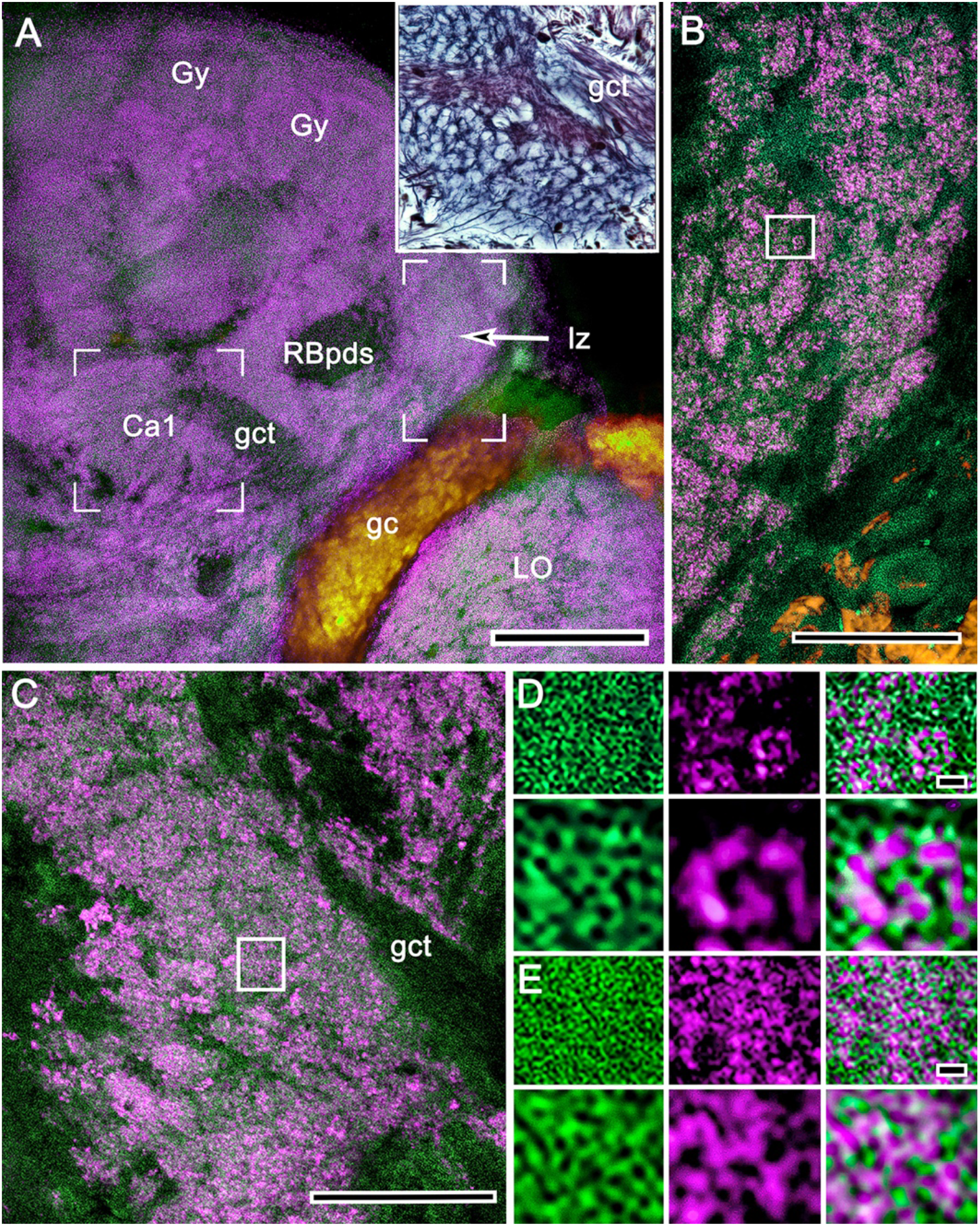
Comparison of synaptic densities associated with the calyces and reniform body. (**A**) Calyx 1 (Ca1), its associated gyri (Gy), the reniform body pedestal (RBpds) and, at this level, its lateral zone (lz) are shown labelled with actin (green)/anti-synapsin (magenta). In A, the open square corresponds to a similar area of silver-stained microglomeruli (upper right inset). The same area is shown enlarged in panel C. The open rectangle in A is enlarged in B to the same scale as panel C. (**B**) Reniform body neuropils comprise synaptic configurations that contribute to large glomerulus-like islets. (**C**) Microglomeruli of the calyces are smaller and more densely packed than those of the reniform body. (**D, E**) Comparison of synapsin/actin contributions to glomerulus-like units of the reniform body (D) and to microglomeruli of the mushroom body calyx (E) further demonstrates major differences regarding the density of their converging presynaptic elements. Other abbreviations: LO, lobula; gc, globuli cells; gct, globuli cell tract. Scale bars, A, 100μm; B,C, 25μm; D, E, 1μm.

